# Psilocybin asymmetrically modulates outcome-based choice and cortical processing under uncertainty

**DOI:** 10.64898/2026.07.16.733837

**Authors:** David Jacobs, Alejandro Torrado Pacheco, Randall Olson, Angela J. Langdon, Bita Moghaddam

**Author notes:** Corresponding Author: Bita Moghaddam 3181 SW Sam Jackson Park Rd #L470 Portland, OR 97239.

## Abstract

Emerging clinical research with psilocybin highlights the value of the psychedelic experience on successful treatment of a multitude of psychiatric disorders. While the psychedelic trip or clinical support procedures cannot be modeled in rodents, neural processes critical to meta-learning and flexible modification of previously learned strategies can be quantified during psilocybin exposure. Here we applied computational modeling and single unit recordings from medial prefrontal cortex to characterize the effect of psilocybin using a value-based probabilistic choice task. Psilocybin improved choice selection in uncertain contexts and shifted reinforcement learning rate bidirectionally, enhancing updating from rewarded, while reducing updating from unrewarded, actions. These behavioral changes coincided with selective shifts in the neural encoding of task features including enhanced neural representation of rewarded, and diminished representation of unrewarded, outcomes. Collectively these data indicate that psilocybin improves choice selection in uncertain contexts by reweighing cortical coding in a way that favors learning from positive new information over prospective actions.

## Introduction

Recent clinical data indicate that psychedelics such as psilocybin hold promise for treating symptoms of mental disorders that are not treated by available pharmacotherapies (Goldberg et al., 2020; Johnson & Griffiths, 2017; Thomas et al., 2017). These data also suggest that psychedelics may work by mechanisms different than classical pharmacotherapy which attribute therapeutic effects to drug action alone, such as the amelioration of anxiety with benzodiazepines. First, the reported clinical efficacy of psychedelics is transdiagnostic, spanning multiple symptoms and disorders ranging from mood disorders, substance use disorders, and pain (Askey et al. 2024; Carhart-Harris et al., 2021; Van der Meer et al., 2023). Second, there is no evidence so far that these drugs work as standalone pharmacology treatment. All the published positive clinical data involve clinical or context-specific intervention during drug administration, and an emerging literature suggests that the acute “experience” during drug exposure predicts therapeutic efficacy (Kerr Gaffney et al., 2026).

Several influential theories propose that the psychedelic experience promotes meta learning and modifies the processing of previously learned associations or held beliefs, leading to a more flexible state (Carhart-Harris & Friston; 2019; Gallimore, 2015; McGovern et al., 2024). Consistent with these theories, animal studies have shown positive effects of psilocybin on cognitive flexibility and different forms of learning (Fisher et al., 2024; Hinchcliffe et al., 2024; Nardou et al., 2023; Torrado-Pacheco et al., 2023). Recent studies have also provided large-scale data for the involvement of prefrontal cortex (PFC) subregions and associated networks (Carhart-Harris et al., 2012; Fuini et al., 2025; Girn et al., 2022) in psychedelic effects. At a mechanistic level, however, animal studies have primarily focused on sustained post-psilocybin effects on synaptic, dendritic, and multi-regional plasticity markers (Shao et al., 2021, 2026; Smausz et al., 2022). Thus, little is known about the experiential impact of psychedelics on neural state dynamics while an organism is engaged in learning and deciding between options (Purple et al., 2025; Rogers et al., 2024).

The current study was undertaken to understand how PFC encoding of decision variables is influenced by psilocybin. We recorded single unit activity from medial tier of the rat PFC (mPFC) during a reward-guided probabilistic decision-making task and applied computational modeling and encoding and decoding analyses. The behavioral task design allowed us to assess adaptive behavior under two conditions: low uncertainty blocks when choice outcome was predictable and high uncertainty blocks when choice outcome was random. Rats learned to adjust their choice dynamically according to shifting probabilities and recent outcomes. Psilocybin enhanced choice variability and increased learning rate for rewarded actions while reducing learning rate for unrewarded actions. At the neural population level, psilocybin produced opposing effects on action outcomes by enhancing rewarded action while attenuating unrewarded action preferring ensembles. These changes were also associated with modified encoding of upcoming choices. These data are consistent with psilocybin enhancing meta-learning and further indicate that it improves choice selection in uncertain contexts by modulating cortical neural dynamics related to favorable outcomes.

## Results

### Psilocybin influences probabilistic choice in an uncertainty dependent manner

Effect of psilocybin on probabilistic decision making was measured using a paradigm adapted from primate and rodent studies (Cheng et al., 2025; Grossman et al., 2022; Sugrue, Corrado, & Newsome 2004). Animals were freely moving in an operant box where they chose between two action options with shifting reward probabilities (**Fig. 1A**). Three unique block types (0.2/0.8, 0.8/0.2, 0.5/0.5) were applied pseudorandomly (**Fig. 1B**). The first choice carried reward probability (p(R)) of 0.2, 0.5, or 0.8, and the second choice had a reward probability of 1-p. This design, which included a “catch-block” where both choice options were reinforced at chance (0.5/0.5), allowed us to assess adaptation to probability shifts on preferred options across two conditions: low uncertainty blocks, where choice outcomes were predictable, and high uncertainty blocks, where outcomes were random. Animals adjusted their choice dynamically according to shifting probabilities and recent outcomes (**Fig. 1C, Supplemental Data Fig. 1A)** and reached stable performance through consecutive daily sessions before receiving psilocybin (see Methods).

**Figure 1.**
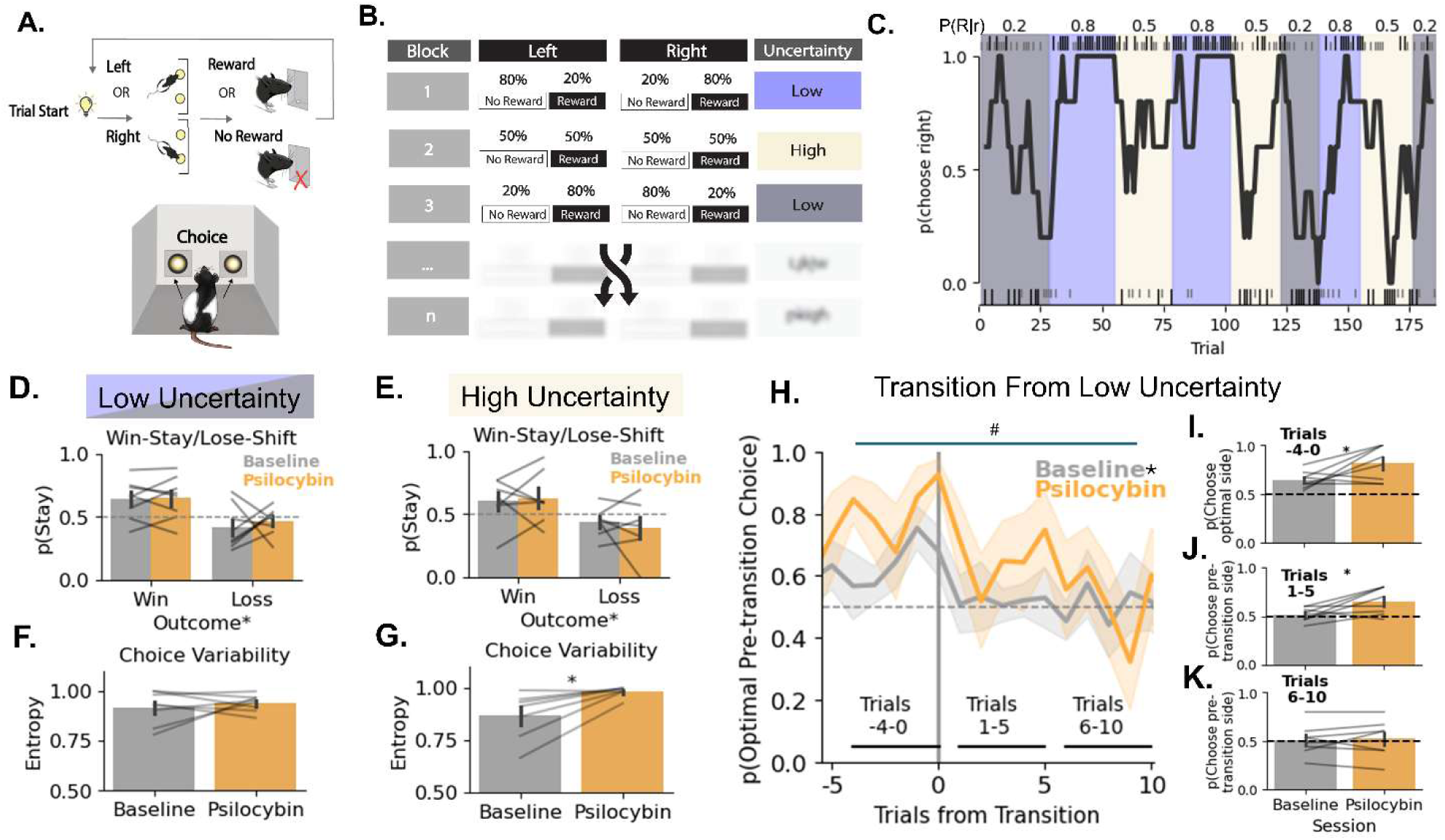
Effects of psilocybin on probabilistic choice behavior. **A.** After a variable amount of time (3-30 sec), a trial began with illumination of the nosepoke light*s*. Animals then made a left or right action (choice), which could immediately result in reward or no reward. Animals were required to make a feeder entry to initiate the next inter-trial interval. **B.** An example session showing the underlying probabilities of reinforcement for left or right choices. Reinforcement probabilities for each action could be one of three combinations. Two blocks were designated as low uncertainty as they contained an optimal choice (e.g. left or right optimal for block 1 and 3 above, respectively). In high uncertainty blocks neither choice was optimal (block 2). Blocks pseudorandomly transitioned to a different block. **C.** Representative choice data (moving 5 trial average) from a subject. Shading represents the underlying block with R=reward, r= choose right. Small hashes at the top and bottom indicate a trial was unrewarded while large hashes indicate a trial was rewarded. **D.** Win-stay/lose-shift for low uncertainty blocks (mean ± SEM, lines indicate individual rats). Animals used recent outcomes to drive choices as evident by outcome sensitive stay probabilities (*p<.05 effect of outcome, RM ANOVA). **E.** Win-stay/lose-shift for high uncertainty blocks (mean ± SEM, lines indicate individual rats). Win-stay/lose shift behaviors were outcome sensitive and unchanged by psilocybin (*p<.05 effect of outcome, RM ANOVA). **F.** Choice variability for low uncertainty blocks (mean ± SEM, lines indicate individual rats). There was no difference between baseline and psilocybin sessions. Note: higher entropy values signify higher variability. **G.** Choice variability for high uncertainty blocks (mean ± SEM, lines indicate individual rats). Psilocybin increased the entropy metric for choice variability (*p<.05, paired t-test). **H.** Transition analysis of choice behavior for low uncertainty blocks (mean ± SEM). Psilocybin enhanced asymptotic (i.e.pre-transition) optimal choice. When contingencies transitioned to lower reinforcement probability for the optimal side, animals adapted by decreasing preference for said side (*significant intercept effect of psilocybin, ^#^p<.05 significant effect of reversal phase, generalized mixed-effects model). **I.** Mean ± SEM and individual subject averages for the pre-transition window (*p=.022, paired t-test one-tailed). **J.** Mean ± SEM and individual subject averages for the 1-5 trial post transition window (p=.036, paired t-test). **K.** Mean ± SEM and individual subject averages for the 6-10 trial post transition window (p=.70, paired t-test). Lines in bar plots indicate individual rats. N=7-8 rats.

The effect of psilocybin (1 mg/kg ip) was compared to animals’ performance on the session before receiving the drug (depicted in all graphs as “baseline”). Psilocybin increased response latency and latency to enter the feeder, and had no generalized effects on accuracy (**Supplemental Data Fig. 1B-D**). However, differences emerged when we compared the performance during low and high uncertainty blocks. Specifically, win-stay/lose-shift and choice variability analyses in low (0.8/0.2) and high (0.5/0.5) uncertainty trials revealed that while outcome related choice strategies remained intact under psilocybin (**Fig. 1D-E**), psilocybin increased variability of choice (entropy) selectively in high, but not low, uncertainty trials (**Fig. 1F-G**). This was an important observation because when outcomes are highly uncertain (which is the case in 0.5/0.5), an increase in choice variability by psilocybin may suggest an increase in adaptive exploration (Padoa-Schioppa, 2013).

We next asked how likely animals were to select the optimal option before the transition to new reinforcement probabilities and then persist in this choice after the block change. To do this we assessed choices across pre (trials −4-0) and post (trials 1-10) transitions using mixed effects models. This analysis indicated that psilocybin treatment was associated with higher asymptotic performance in trials leading up to transitions out of low uncertainty, whereas it had no interactive effects on post-transition parameters (**Fig. 1E, Supplemental Data Fig. 1F; Statistics Supplement**) and no comparable effect for transitions out of high uncertainty (**Supplemental Data Fig. 1F**). Analysis of binned individual responses over 5 trial windows supported the pre-transition improvement for the majority of psilocybin-treatment animals (**Fig. 1I**). While, after transition, most psilocybin-treated animals had an initial larger probability of maintaining the previously preferred option (**Fig. 1J**), they returned to baseline levels after 5 trials (**Fig. 1K).** These findings suggest that acute psilocybin improves choice selection in low-uncertainty contexts.

Of note, a subset of animals (5 of 8) received saline injections in sessions other than the psilocybin-injection session and these injections did not influence the behavior (**Supplemental Data Fig. 1G**). Because of unanticipated technical issues, not all animals received a saline injection. The subsequent analyses, therefore, focus on comparing the effects of psilocybin on animals’ baseline.

### Psilocybin attenuates choice bias and asymmetrically shifts outcome specific updating

To ask how psilocybin produces its effect on dynamic choice options, we fit 5 reinforcement learning models with varying complexity to behavioral data (see Methods), and modeled psilocybin treatment as a group level effect on parameter estimates similar to human psychedelic probabilistic learning studies (Kanen et al., 2022). All models used recent outcomes to update values for each choice option, with probabilistic action selection based on these values. Our preferred model (**Supplemental Data Fig. 2A**) included 4 parameters of interest: separate update rates for rewarded (η+) and unrewarded (η-) outcomes, a bias parameter capturing value independent preference for one side, and an inverse temperature parameter capturing the degree to which choices were driven by learned values (**Fig. 2A**).

**Figure 2.**
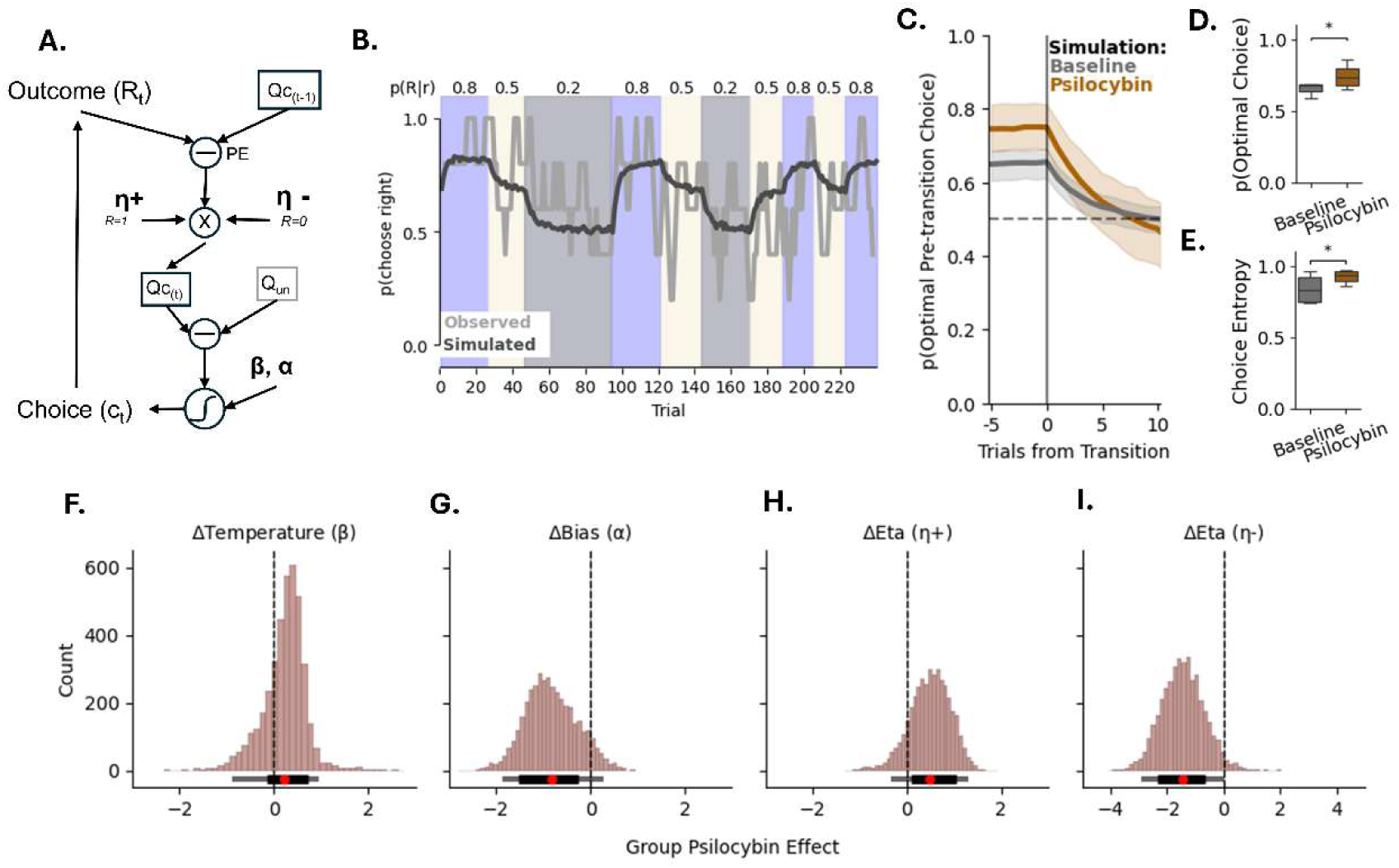
Computational modeling of choice behavior indicates psilocybin shifts learning mechanisms. **A.** Schematic of the reinforcement learning process with outcome dependent learning rates (η+/-), a value independent bias parameter (α), and an inverse temperature (β) to influence choice policy via Q learning. Choices (c_t_) produce an outcome (R_t_) which is compared to the prior Q value for a given chosen side (Qc_(t-1)_) to form a prediction error (PE). Learning rates then control how much the PE is used to update Qc. The Q value for the chosen and unchosen (Q_un_) options are transformed into choice probabilities via a logistic function. **B.** Representative observed choice behavior (5 trial average, light grey) and simulated model predictions for that subject (dark grey). Shading indicated probability of reward given a right choice. **C.** Simulated choice averages (mean and 95% CI) around transitions in reinforcement probability from low uncertainty for baseline and psilocybin sessions. **D.** Boxplots of model simulations (median and IQR, whiskers indicate 5-95 percentile) for asymptotic optimal choice in low uncertainty blocks (trials −4-0). The model recovered increases in optimal choice seen after psilocybin treatment (*p=.025, one-tailed t-test). **E.** Boxplots of model simulations (median and IQR, whiskers indicate 5-95 percentile) for choice entropy in high uncertainty blocks. The model recovered increases in choice entropy seen with psilocybin treatment (* p<.01, one-tailed t-test). **F-I.** Posterior parameter distributions for the group level effect of psilocybin (n=4000 estimates from 4 chains). Below the distribution: red circles indicate the mean, thick black bars indicate 80% highest density interval (HDI) and thinner black bars indicate 95% HDI. **F.** Temperature was not affected by psilocybin (0∊75%HDI). **G.** Bias was attenuated by psilocybin (0∉80%HDI). **H-I.** Psilocybin shifted learning rates by increasing rewarded learning rate (0∉80%HDI) and decreasing unrewarded learning rate (0∉90%HDI).

Two checks confirmed the model captured behavior. First, simulating subject data from best fit parameters recapitulated choice behavior in the task (**Fig. 2B**), including the dynamic choice preferences during transitions from low uncertainty (**Fig. 2C**). Second, applying the fitted psilocybin effect onto each of the parameters in the model recovered the key behavioral signatures of psilocybin treatment: increases in optimal choice during low uncertainty (**Fig. 2D**) and increases in choice variability during high uncertainty (**Fig. 2E, Statistics Supplement**).

Posterior estimates for psilocybin effects indicated there was no evidence for an effect of psilocybin on temperature (0 ∊ 75% HDI, dBF< 3.2, **Fig. 2F**) but moderate evidence for decreases in bias (0 ∉ 80% HDI, dBF = 11, **Fig. 2G**). The strongest psilocybin effects were on updating mechanisms. Psilocybin moderately increased learning rate for rewarded and strongly decreased learning rate for unrewarded outcomes (Δη+ = 0 ∉ 80% HDI, dBF = 7.1, **Fig. 2H**; Δη-= 0 ∉ 90% HDI, dBF = 31, **Fig. 2I**). These findings indicate that psilocybin reshapes choice and update mechanisms under uncertainty in two ways: by attenuating value-independent choice bias and asymmetrically reweighting how rewarded versus unrewarded outcomes update choice values.

### Psilocybin shifts neuronal representation of outcome in the mPFC

During the behavioral task described above, single unit spike activity from mPFC was recorded (Baseline n=207 units, Psilocybin n=222 units; **Supplemental Data Fig. 3A-D**). Guided by the outcome of reinforcement learning analyses described above, we analyzed the population activity trajectory for rewarded and unrewarded trials. Action-outcome activity trajectories were well described by a third order polynomial (**Supplemental Data Fig. 3E-F; Statistics Supplement**). Psilocybin altered these trajectories resulting in elevated post-outcome activity in rewarded trials (**Fig. 3A**) and attenuated activity in unrewarded trials (**Fig. 3B**) relative to baseline.

**Figure 3.**
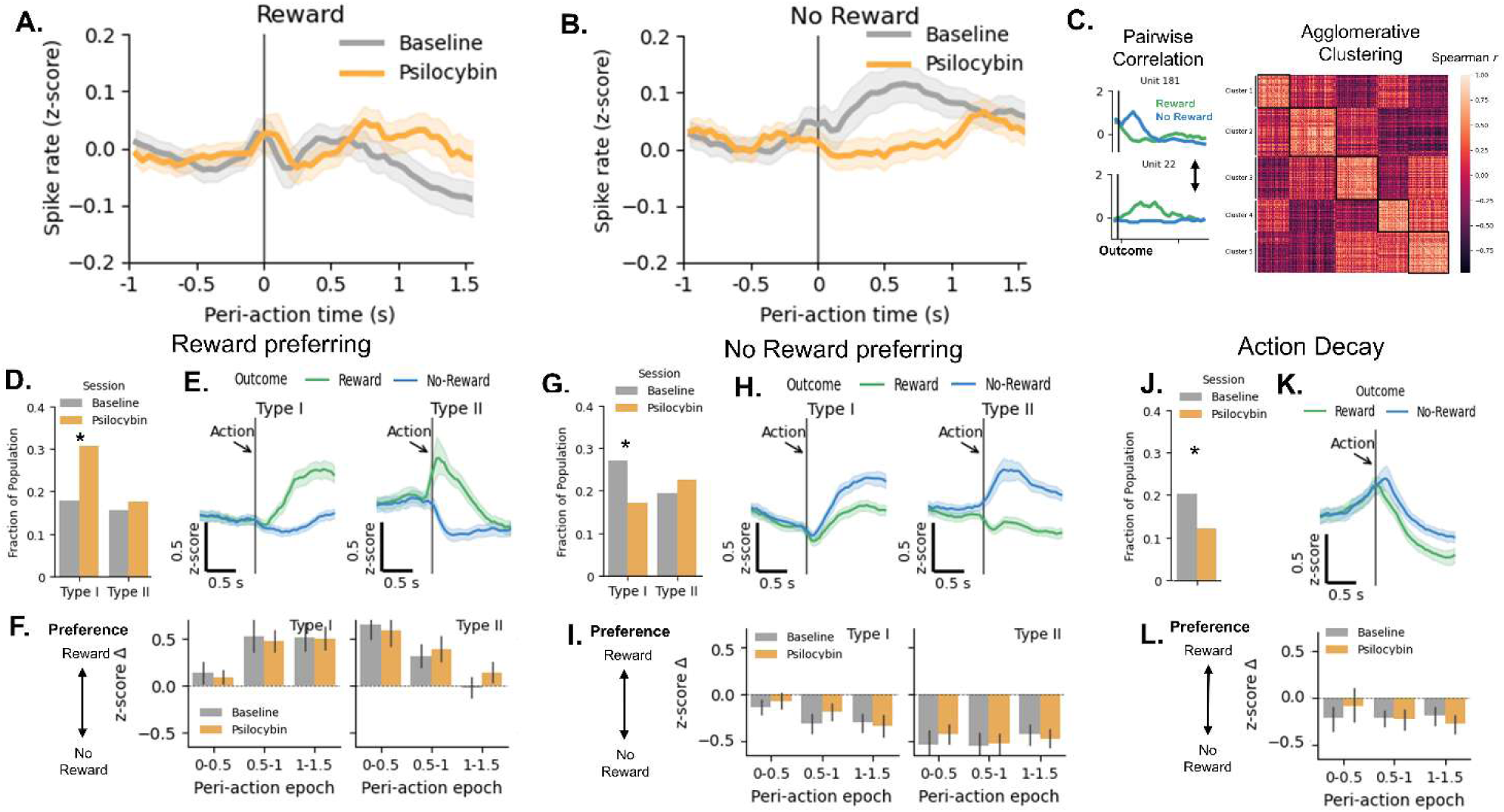
Psilocybin bidirectionally shifts population and subcluster related activity to reward and no reward outcomes. **A-B.** Population averaged firing rate during the peri-action epoch for trials which resulted in reward or no reward (mean ± SEM). Psilocybin altered response trajectories by elevating the length of reward response and inhibiting no reward related response relative to baseline (n=207-222 units from 7-8 rats; mixed effects model interactions p < .02). **C.** Schematic demonstrating reward and no reward responses of two example units. These signals were correlated between each unit pair and subjected to clustering analysis based on reward and no reward response profiles (see methods). **Right** - Unit by unit correlation matrix sorted by the results of agglomerative cluster analysis. **D.** Two subtypes of reward-preferring neuronal clusters were found based on their temporal pattern of activation (depicted as Type I and Type II). Psilocybin selectively increased the relative proportion of Type I units (* p=.01, Fisher exact test, Benjamin-Hochberg corrected). **E.** Mean and 95% CI for peri-action activity of each subtype (n=71-105 units). **F.** Mean and 95% CI relative change between reward and no reward response shows preference for reward which was not influenced by psilocybin. **G.** Two subtypes of no reward preferring neuronal clusters were found, and psilocybin selectively decreased the relative proportion of Type I units (* p=.03 Fisher exact test, Benjamin-Hochberg corrected). **H.** Mean and 95% CI for peri-action activity of each subtype (n=90-94 units). **I.** Mean and 95% CI relative change between reward and no reward response shows preference for no reward, the magnitude of which was not influenced by psilocybin. **J.** A final cluster of neurons showed rapid decay in activity immediately at action/outcome, and this cluster was less prevalent during acute psilocybin (* p=.04, Fisher exact test, Benjamin-Hochberger corrected). **K.** Mean and 95% CI for peri-action activity of the final cluster (n=69 units). **L.** While decreases were observed in activity for rewarded and unrewarded trials, preference scores indicate that magnitude of this decrease was lower for unrewarded trials (mean and 95% CI).

PFC neurons are known to show heterogenous responses to task variables including outcomes (Del Arco et al., 2017; Le Merre et al., 2026), which are not captured by overall population averages. To better understand post-outcome signaling changes, we calculated pairwise correlations between units (**Supplemental Data Fig. 4A**) and utilized hierarchical clustering to identify distinct subclusters of neurons in the mPFC based on response to rewarded and unrewarded trials (**Fig. 3C**). Five clusters robust to leave-one-out stability analysis (mean adjusted Rand index =.84; **Supplemental Data Fig. 4B-C**) were identified. These clusters were classified by outcome preferences and activity patterns. There were two clusters of neurons which showed post-outcome activity increases that were higher for rewarded outcomes (Reward preferring). These neurons had distinct temporal patterns, showing reward-related increases immediately at action or later in the action/outcome epoch and are depicted as Type I and Type II, respectively (**Fig. 3D-E**). Psilocybin increased Type I, but not Type II, neural representation as indicated by increases in overall proportion of the population. Calculating preference scores for these neurons indicated that the magnitude of preference for reward was not affected by psilocybin (**Fig. 3F**). Two other clusters of neurons also showed increases in post-outcome activity, but preferred unrewarded outcomes (No Reward preferring). Like the Reward preferring clusters, No Reward preferring clusters had unique temporal patterns of response relative to action/outcome delivery (Type I and II; **Fig. 3G-H**). Here, psilocybin selectively attenuated representation of the Type I subgroup without affecting the magnitude of preference for unrewarded outcomes (**Fig. 3G-I**). The final cluster displayed a ramping of activity pre-action and decayed at outcome. This group decreased in representation with psilocybin treatment (**Fig. 3J-K**). Overall, decay was more modest following unrewarded outcome in these neurons, and this preference was similar across baseline and psilocybin sessions (**Fig. 3L**). These findings suggest that psilocybin bidirectionally modifies the outcome representation in the PFC: it enhances representation of rewarded actions and attenuates representation of unrewarded actions.

### Psilocybin attenuates choice-related encoding in the mPFC

Having found that psilocybin reshapes outcome representations, we next asked whether its effects extended to other task variables, most importantly the encoding of choice-related signals. We fit regressions to single-neuron activity to determine which aspects of the task were encoded across baseline and psilocybin sessions. Overall, a large portion of neurons encoded at least one task variable in baseline and psilocybin sessions (Baseline: 72% of units, Psilocybin: 73% of units), and mixed encoding was observed regardless of treatment (**Supplemental Data Fig. 5A-B**). Choice direction encoding (regardless of outcome) was observed in 36-39% of neurons and encoding was highest in the pre-action period (representative unit in **Fig. 4A**). Psilocybin attenuated this response when compared to baseline during the pre-action period (**Fig. 4B; Statistics Supplement; Supplemental Data Fig. 5A-B**). mPFC neurons also encoded trial outcome (representative unit in **Fig. 4C; Supplemental Data Fig. 5A-B**). Outcome encoding peaked immediately after action (i.e. at outcome delivery) but was not significantly influenced by psilocybin (**Fig. 4D**). Notably, conditional encoding of outcome as a function of choice peaked after outcome delivery and was attenuated by psilocybin (**Fig. 4E-F**). This effect was only significant in the mixed effects analysis, suggesting it reflects a diffuse reduction in encoding magnitude rather than changes in precise timepoints. While prior choice encoding was minimal (**Supplemental Data Fig. 5C-D**), high levels of prior outcome encoding were seen in the ITI (**Supplemental Data Fig. 5E**). This prior outcome encoding continued into the peri-action epoch and was modestly enhanced by psilocybin, albeit not significantly after post-hoc correction (**Fig. 4G-H**). Together these findings demonstrate that psilocybin disrupts choice signals and choice-related outcome encoding while preserving responses to recent outcome history. This raised a more specific question: because choice signals can be driven by underlying value computations, was the attenuation of choice encoding accompanied by a parallel disruption of value signaling?

**Figure 4.**
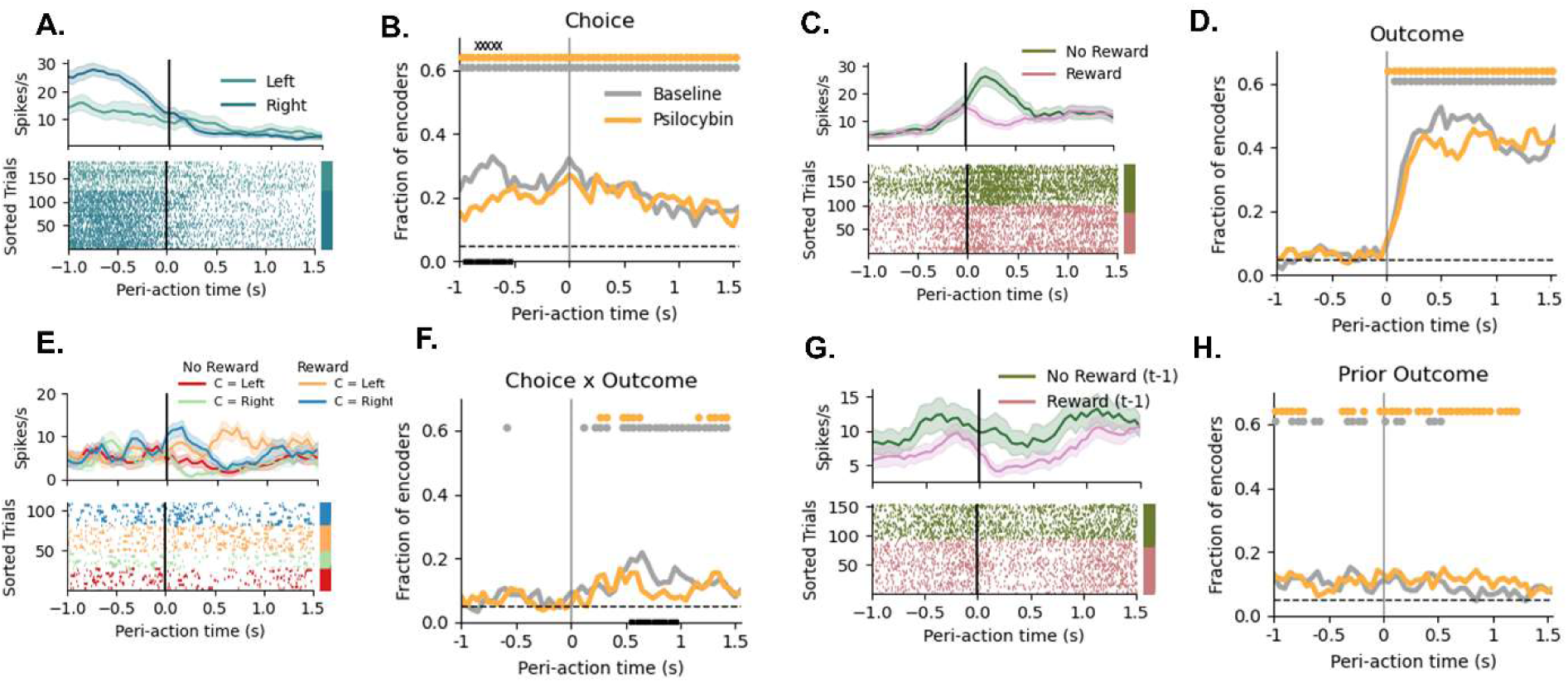
Psilocybin attenuates choice related encoding at multiple timescales. **A.** Mean spike rate and raster plot for a representative single unit which encodes choice direction (left/right). **B.** Fraction of task encoders showing significant choice encoding at each time bin over the peri-action period. Psilocybin attenuated encoding prior to the action being executed. **C.** Mean spike rate and raster plot for a representative single unit which encodes outcome. **D.** Fraction of task encoders showing significant outcome encoding at each time bin over the peri-action period. **E.** Mean spike rate and raster plot for a representative single unit which encodes outcome differentially based on choice. **F.** Fraction of task encoders showing significant choice x outcome encoding at each time bin over the peri-action period. Psilocybin attenuated encoding following outcome delivery. **G.** Mean spike rate and raster plot for a representative single unit which encodes the outcome from the prior trial. **H.** Fraction of task encoders showing significant prior outcome encoding at each time bin over the peri-action period. Colored circles above plots indicate significant binomial test (p<.01 Benjamin-Hochberg corrected) for baseline or psilocybin. Black ‘x’ above plots indicates significant cluster corrected chi^2^ test between baseline and psilocybin. Solid black bars on x-axis indicate significant session difference for that 0.5 sec time chunk (GLM with Benjamin-Hochberg corrected post-hoc contrast).

To address this, we extracted relative and total value signals from our computational modeling of choice behavior and ran an additional choice-value regression. Decision value signals have been implicated in biasing action selection and tracking reward richness in the environment (Bari et al., 2019; Hamid et al., 2014; Ito & Doya, 2009; Niv et al., 2007), and would be a natural locus for psilocybin’s effect on choice if value computations themselves were disrupted. Similar proportions which encoded one or both of the value signals were observed (Baseline: 20% of units, Psilocybin: 24% of units; **Supplemental Data Fig. 6A-B**). Relative value signals track optimal choice and may guide action selection (**Fig. 5A**). Consistent with previous studies (Bari et al., 2019), we found neurons which modulated activity monotonically with relative value (representative unit in **Fig. 5B**). Relative value encoding was sustained over the action/outcome epoch and encoding was observed under psilocybin. There was a trend for attenuation of relative value encoding in the psilocybin session but this effect was not significant after post-hoc correction (**Fig. 5C; Statistics Supplement**). This pattern may be expected given that relative value is thought to influence selection of choice, and that both choice and choice-by-outcome encoding were similarly attenuated in our task variable model. Unlike relative value, total value signals follow unique dynamics related to reward history and updating that are believed to support state value signals (**Fig. 5D**). While total value neurons were observed (**Fig. 5E**), they showed a narrower window of encoding above chance and were not significantly influenced by psilocybin (**Fig. 5F**). Collectively these analyses suggest that choice-related value signaling in the mPFC is largely maintained during decision making under psilocybin. The attenuation of choice encoding therefore appears not to reflect a disruption of upstream value computations, prompting us to ask *when* in the trial the choice signal is degraded.

**Figure 5.**
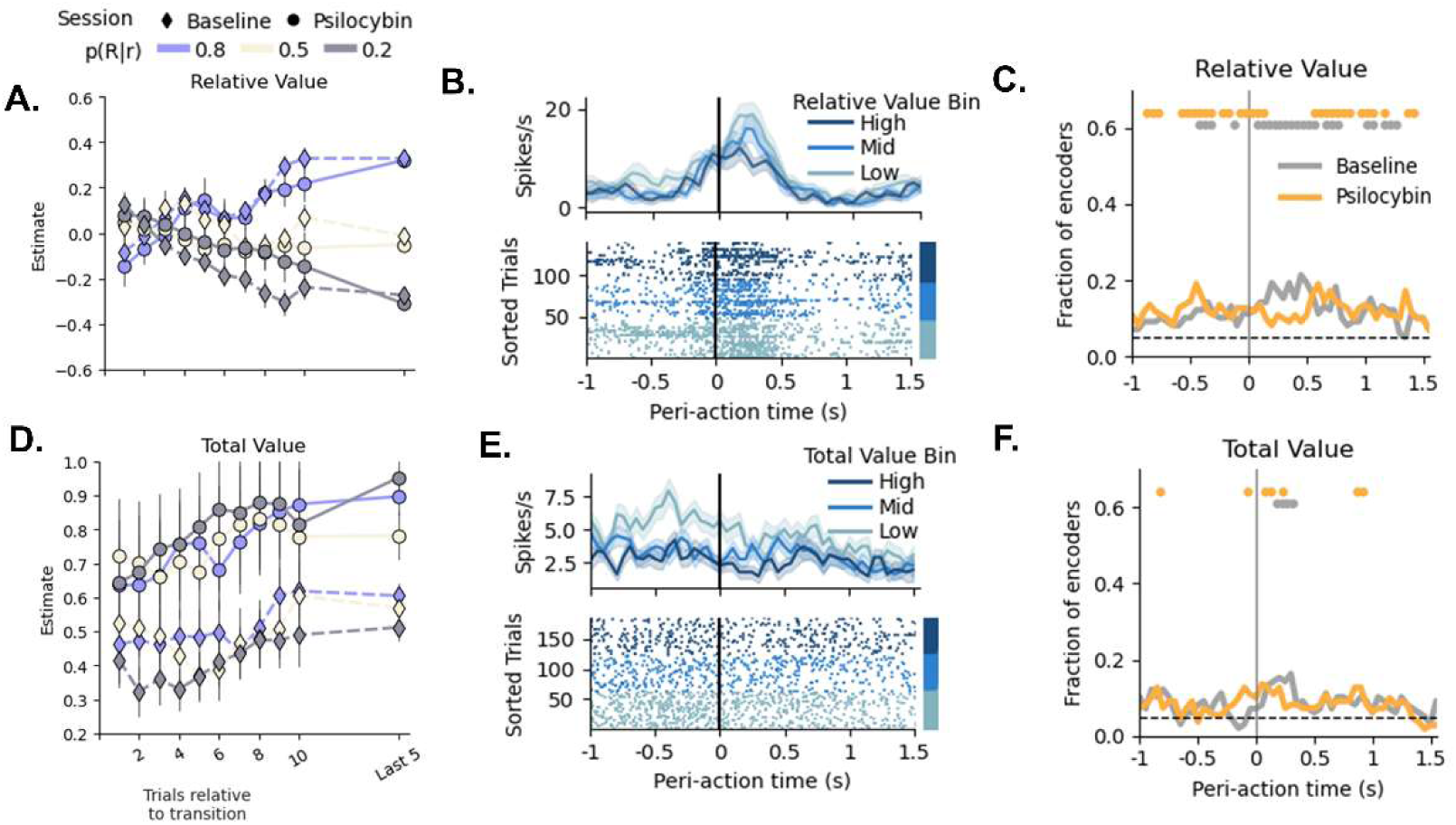
Encoding of total and relative value signals during acute psilocybin. **A.** Mean and SEM of relative value estimates in the first 10 trials of a block and the last 5 trials of the block. Data are color coded by block type and symbols reflect psilocybin (circles) or baseline (diamond) session. **B.** Mean spike rate and raster plot for representative single unit with relative value encoding. For visualization, data are colored by terciles of value estimates. **C.** Fraction of choice-value encoders showing significant relative value encoding at each time bin over the peri-action period. **D.** Mean and SEM of total value estimates in the first 10 trials of a block and the last 5 trials of the block. Data are color coded by block and symbols reflect psilocybin (circles) or baseline (diamond) session. **E.** Mean spike rate and raster plot for representative single unit with total value encoding. For visualization, data are colored by terciles of value estimates. **F.** Fraction of choice-value encoders showing significant total value encoding at each time bin over the peri-action period. Colored circles above plots depict significant binomial test (p<.01 Benjamin-Hochberg corrected) for baseline or psilocybin.

### Psilocybin changes coding for choice signals before, but not after, action execution

To localize this effect in time, we used linear decoding to ask how populations of mPFC neurons predict upcoming choice direction in the action/outcome epoch. We found that mPFC pseudo-populations show above chance decoding of upcoming choices in the pre-action period (**Fig. 6A; Statistics Supplement**). Psilocybin decreased decoding across pseudo-populations and did not achieve the same level of decoding even with the full population. However, analysis of decoding accuracy across the epoch revealed that choice decoding returned to baseline levels after action execution (**Fig. 6B, Supplemental Data Fig. 7A**), suggesting that psilocybin influences upcoming but not retrospective chosen action related signals. Next, we asked if some of the choice decoding attenuation could be explained by subpopulations identified by cluster analysis. We re-ran decoding analysis while leaving out specific subclusters (i.e. those in **Fig. 3**) and calculated the change in predictive accuracy from the full population. This analysis revealed that action-decay neurons, which were attenuated by psilocybin (**Fig. 3J-K**), had the largest influence on choice decoder accuracy across sessions. Although the magnitude of action-decay-related losses in decoding accuracy was modest, these losses were larger than those produced by leaving out the other psilocybin-affected subpopulations (Reward Type I and No Reward Type I; **Fig. 6C**). Thus, psilocybin’s reweighting of the mPFC population may shift the network toward a state that prioritizes reward processing at the expense of ongoing action-related activity. Consistent with this interpretation, outcome decoding was largely unaffected by psilocybin at higher pseudo-population sizes (**Supplemental Data Fig. 7B-D**).

**Figure 6.**
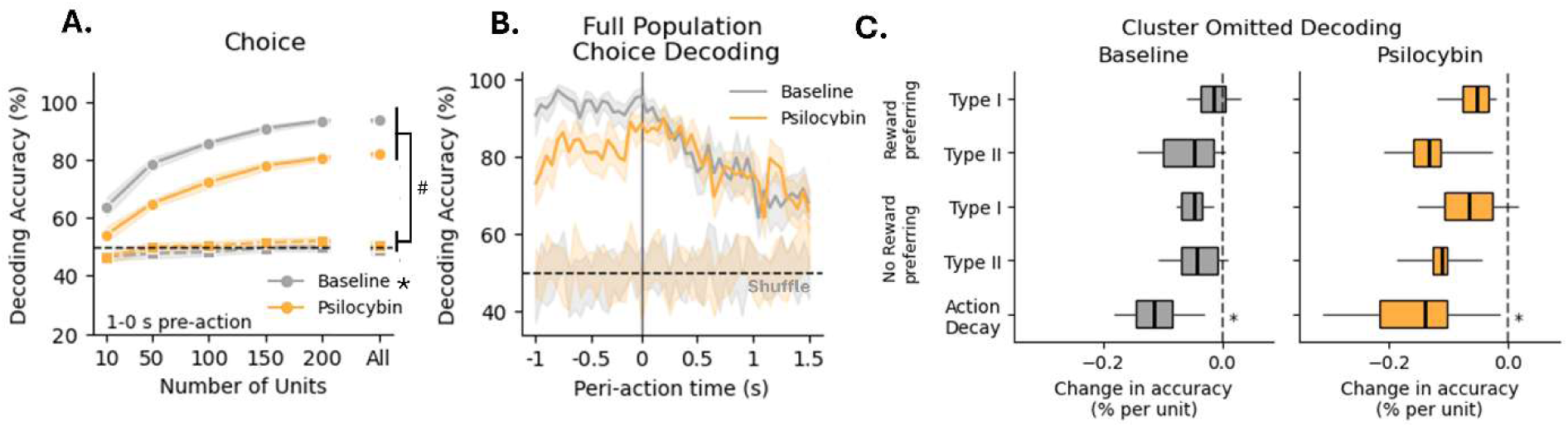
Psilocybin attenuates decoding of choice in the peri-action period. **A.** Decoding choice (chosen side) before action execution was attenuated by psilocybin across a range of pseudo-population sizes (mean ± 95%CI, *p<.05 main effect of psilocybin). Squares near the 50% chance line are the results of shuffle analysis (^#^p<.05 versus shuffle). **B.** Full population decoding (mean ± 95% CI, n = 20 runs with 207-222 units) demonstrated that decoding performance for choice was highest before and at action execution in baseline sessions. Decoding performance was highest at action execution in psilocybin sessions. Lighter shading indicates mean ± 95%CI for shuffle runs. **C.** Box plots (median ± whiskers for the 5-95 percentile) showing effects of removing a specific cluster (see Figure 3) from choice decoding analysis during the pre-action period (−1-0 s). The largest losses in accuracy were observed when decay neurons were ablated (*p<.05 action decay vs each other cluster, Benjamin-Hochberg corrected).

## Discussion

Therapeutic effects of psychedelic-assisted therapy are commonly attributed to meta-learning and integration of new learning and modulation of how previously learned rules and beliefs (priors) guide behavior. These assumptions are consistent with data showing that psilocybin increases behavioral flexibility in clinical populations (Doss et al., 2021; Sloshower et al., 2024) and in animal models (Conn et al. 2024; Torrado-Pacheco et al.; 2023; Woodburn et al., 2024). Neurocomputational mechanisms that may guide these behavioral effects remain elusive. While many large-scale basic and clinical studies have identified prefrontal cortical networks as key responders to psychedelic exposure (Davoudian 2023; Jiang et al., 2025; Siegel et al., 2024), these studies have primarily focused on static molecular and structural consequences of psilocybin (Girn et al., 2026; Kwan et al., 2022; Smausz et al., 2022). Here we sought to understand the dynamic impact of psilocybin on cortical processing during execution of outcome-based flexible learning. We thus examined the effect of psilocybin on PFC neuronal activity while rats were engaged in a probabilistic choice task along with computational modeling of the behavioral effects. We find that psilocybin improves choice during low uncertainty trials while increasing exploration during high uncertainty trials. These behavioral effects of psilocybin were associated with an attenuation of choice biases and preferential activation of PFC neuronal responses to rewarded choices.

### Psilocybin modulates probabilistic choice behavior and reinforcement learning rate in favor of positive outcomes

Psilocybin shifted probabilistic choice in an uncertainty-specific manner. When uncertainty was low, psilocybin improved asymptotic performance, consistent with facilitation of behavioral flexibility reported in humans and rats, while having no effect on other behavioral strategies such as win-stay/lose-shift (Casanova et al., 2024; Doss et al., 2021; Rogers et al., 2025; Torrado-Pacheco et al., 2023; Woodburn et al., 2024). These results are consistent with the idea that acute psilocybin improves meta-learning by flexible adaptation of choice when uncertainty levels are dynamic.

Choice behavior may be influenced by multiple processes related to choice biases, value estimates, uncertainty, and outcome updating, many of which may be modulated by serotoninergic signaling (Cohen et al., 2015; Colwell et al., 2024; den Ouden et al., 2013; Grossman et al., 2022; Harkin et al., 2025; Lottem et al., 2018; Matias et al., 2017). To better understand how psilocybin influences these decision processes in our task, we used reinforcement learning models. We found that psilocybin produces outcome-specific bidirectional effects on learning rate: increasing updating after rewarded outcome, while decreasing updating after unrewarded outcome. These data support the notion that psilocybin reweighs update mechanisms in favor of positive feedback while countering negative feedback. These results are clinically relevant observations and consistent with recent human studies reporting that LSD, another serotonergic psychedelic, increases learning rates of rewarded choices in a probabilistic reversal learning task (Kanen et al., 2022). Our results are also supported by recent studies showing that psilocybin reduces bias from negative outcomes in rats (Fisher et al., 2024; Hinchcliffe et al., 2024; Jacobs et al., 2024) and attenuates negative affective biases in humans (Martens et al., 2025).

The effects of psilocybin in our task were not solely due to shifts in learning but also related to choice processes under uncertainty. In particular, the value-independent choice bias (i.e. the side bias) parameter was decreased by psilocybin. This parameter captures the propensity to choose a particular side option, which can have a larger impact when options have similar values relative to one another, a dynamic that is common when uncertainty is high. Using entropy-related analysis to quantify variability of choice (Woo et al., 2023), we found that psilocybin increased choice entropy only under high uncertainty where values for each action are similar and exploring both options is useful for detecting when the contingencies switch. Collectively, these findings suggest that psilocybin promotes exploratory behavioral patterns, like that observed with LSD (Casanova et al., 2024; Conn et al., 2024; Kanen et al., 2022; Muller et al., 2025).

### Psilocybin reorganizes mPFC neuronal populations and encoding properties

A large literature has reported on the PFC encoding of various aspects of action selection and decision processes (Angelaki et al., 2025; Domenech & Koechlin, 2015; Findling et al., 2025; Glascher et al., 2009; Soltani & Izquierdo, 2019; Steinmetz et al., 2019). In similar probabilistic choice tasks in rats and primates, PFC shows high levels of activity during choice, outcome, and value estimates (Bari et al., 2019; Choi et al., 2023; Tsutsui et al., 2016). The PFC receives extensive serotonergic innervation from raphe nuclei and contains 5-HT2 receptors (Aghajanian et al., 1970; Shin et al., 2021; Shao et al., 2025). Relevant to the current study, activity of serotonin raphe neurons correlates with uncertainty levels in a similar task (Grossman et al., 2022).

Consistent with this literature, we found that groups of mPFC neurons responded to positive (choice leading to reward) or negative (choice leading to no reward) outcomes. Psilocybin modified this representation in an outcome specific manner: it enhanced the expression of positive outcome clusters while inhibiting the expression of negative outcome clusters. This observation coincided with the directions of update related parameters predicted by the reinforcement learning model. At the single neuron level, outcome-related encoding was maintained as was the ability to decode outcomes from population activity, suggesting the capacity to discriminate between outcomes was not disrupted by psilocybin.

Unlike outcome encoding, choice encoding was attenuated by psilocybin at the single neuron level. The selectivity of this effect was supported by a modest, albeit non-significant attenuation of ‘relative value’ signals related to choices, and a preservation of motivational ‘total value’ signals (Bari et al., 2019; Cheng et al., 2025). This may be explained by the ensemble shifts outlined in the cluster analysis. In addition to outcome preferring subclusters, a subcluster of neurons which increased activity pre-action and decayed after action (choice) were attenuated by psilocybin. Using decoding analysis, we found that choice decoding accuracy was only attenuated in the pre-action period when it also had the largest influence on choice decoding. These findings suggest that psilocybin may shift mPFC ensemble activity away from prospective choice processing in favor of outcome sensitivity.

Recent work using single unit recordings show unidirectional shifts in spontaneous cortical activity and mPFC and ACC neurons by psilocybin, or the pharmacologically related drug ±DOI (Golden & Chadderton, 2022; Purple et al., 2025; Wood et al. 2012). These effects were, however, observed preferentially during resting, off-task behavior. In rats performing a fear extinction task, single neuron calcium imaging has demonstrated retrosplenial cortical activity is bidirectionally modulated at the ensemble level (Rogers et al., 2025). Similarly, our findings here indicate mPFC subpopulations and encoding patterns can be increased or attenuated by psilocybin in an event-specific manner. This highlights the importance of studying the effects of psilocybin on PFC activity (as well as other regions) when there is dynamic behavioral or contextual engagement rather than during resting or static/postmortem conditions.

### Implications for therapeutic mechanism and related theories

Psilocybin’s therapeutic potential is bolstered by its apparent transdiagnostic effects, targeting common mechanisms across mental health disorders such as depression, anxiety disorders, and substance use (Kelly et al., 2021). These disorders commonly share maladaptive learning and sensitivity to negative feedback, and cognitive rigidity (Balconi et al., 2014; Eshel et al., 2010; Meiran et al., 2011; Pizzagalli, 2011). Furthermore, deficits in reinforcement learning parameters have been documented in patients with these disorders (Brown et al., 2021; Gueguen et al., 2021; Langley et al., 2025; Maia & Frank, 2011; Reinen et al., 2021). In this framework, psilocybin may increase reward learning rates to allow patients to more rapidly reweight positive associations which drive behavior, while reduced non-reward learning rates and bias could facilitate disengagement from negative thought patterns. Thus, modulation of learning parameters provides one potential mechanism by which psychedelics could produce a state to counter the symptomology of rigid cognitive and motivational biases, though it’s important to note that this state could be detrimental in the wrong context (e.g. uncontrolled therapeutic settings; McGovern et al., 2024). While our data do not address whether psychedelics produce long-lasting changes, acute effects are important for treatment outcomes (Kerr Gaffney et al., 2026; Wulff et al., 2023). A transient effect on processing and cortical activity may create a window of enhanced plasticity which would be favorable for therapeutic intervention.

Our data may also have implications for recent theories on how acute psychedelic exposure alters brain processing by “relaxing” prior information and decreasing top-down constraint (Carhart-Harris & Friston, 2019). Related models contend psychedelic effects on memory-associated regions such as the hippocampus also allow higher flexibility in other regions such as the cortex (McGovern et al., 2024). This interplay of relaxed constraints and increased weighting on novel input facilitates updating of beliefs about the environment. Some aspects of our findings are consistent with these theories. For example, the choice encoding shifts we observe with psilocybin may reflect a decrease in prior biases allowing a ‘freer’ landscape for choice, especially under high uncertainty.

Furthermore, changes in outcome-specific representation in PFC may support shifts in updating and scaling of outcome-based prediction errors.

## Conclusions

We find that psilocybin improves choice selection in uncertain contexts by shifting mPFC encoding in an outcome-selective manner. Medial PFC subregions are key nodes within action and outcome processing, commonly involved in top-down control. During a probabilistic choice task, psilocybin increased choice variability and learning rate for rewarded actions while reducing learning rate for unrewarded actions. At single unit and population levels, it caused a dynamic shift in mPFC activity by enhancing reward preferring ensembles, attenuating unrewarded action preferring ensembles and reducing encoding of upcoming choices. These findings support the idea that psilocybin reweighs cortical coding in a way that favors learning from positive new information over prospective action signaling.

## Methods

### Subjects

Twelve-week-old Long-Evans rats (n = 8, 4 of each sex) were purchased from Charles River Laboratories (Wilmington, MA). Rats were pair-housed in same-sex pairs in a room with controlled humidity and temperature under a 12h reverse light/dark cycle. Behavioral testing was conducted during the active dark phase. Rats were allowed to acclimate for one week, after which food restriction began. Rats were fed 11-13g of chow daily until they reached 85-90% of their free-feeding weight. Rats were also handled daily during this period to habituate them to manipulation by experimenters. Once rats reached their target weight, behavioral training began. Rats were weighed daily during behavioral testing and feeding amounts adjusted to maintain the restricted weight, allowing for an increase of 2-5g of weight per week until the rats reached 18 weeks of age.

### Behavioral Apparatus and Training

Training in the behavioral task took place in sound-insulated operant chambers (Coulbourn Instruments) containing a food trough on one wall and two nose-poke ports on the opposite wall. Rats were first acclimated to the operant chambers for two 20-minute sessions occurring on consecutive days, during which sucrose pellets (45 mg, Bio-Serv, Frenchtown, NJ) delivered at random times (5-20 second intervals) in conjunction with illumination of the food delivery port to signal reward availability. Animals were then trained to respond to one of the nose-poke ports (counter-balanced across animals) on a fixed-ratio 1 (FR1) schedule. On each trial both ports were illuminated. Responses at the rewarded side resulted in illumination of the food trough and delivery of one sucrose pellet. Responses on the unrewarded side resulted in illumination of the food trough without concomitant reward delivery. A new trial did not begin until animals nose-poked in the food trough. This phase of the training lasted until animals obtained 50 rewards within a 60-minute period in two consecutive sessions. Sessions in the FR1 phase were conducted daily and the rewarded side alternated in each daily session, to ensure rats learned to respond to both ports. Following successful acquisition of FR1 training (4-6 sessions per animal), training in the probabilistic choice task began.

### Probabilistic Choice Task

The probabilistic choice task used in this study was adapted from a previously published paper using head-fixed mice (Grossman et al. 2022). The task was organized in three alternating blocks, for which the probability of rewards were: [0.8, 0.2]; [0.2, 0.8]; or [0.5, 0.5]; where the numbers correspond to the probability of reward on each trial at the nose-poke ports ([Left, Right]). Blocks were presented in pseudorandom order, where each sequence of 3 blocks was a shuffled presentation of the 3 possible reward blocks. This ensures that the same reward probabilities did not occur in more than 2 consecutive blocks. Block lengths were chosen from a uniform distribution of 15-30 trials. The task ended after 1 hour. Training consisted of running the animals in the task daily. Animals were run on the task until the average performance across the cohort was above chance (i.e. above 50%) for 3 consecutive sessions.

### Drugs

Psilocybin was a gift from the National Institute on Drug Abuse (NIDA) or from the Usona Institute’s Investigational Drug & Material Supply Program. All injections were dissolved in sterile saline were and administered 15 min before the testing session at a dose of 1 mg/kg.

### Electrode implantations

#### Custom built microdrives

Rats were implanted with Custom 64 channel microdrives, which were used for recording single-units in the medial prefrontal cortex (32 channels per hemisphere). Microdrives were constructed as by using two 32-channel electrode interface boards (EIB-36-Narrow-PTB, Neuralynx, Inc., Bozeman, MT, USA) threaded with tungsten wire (12 µm diameter; California Fine Wire Company, Grover Beach, CA, USA), and securing the wires in the board with gold pins (Neuralynx, Inc., Bozeman, MT, USA). Each 32-channel bundle of tungsten wire was threaded through polyimide microtubing (MicroLumen, 160-I) for stability. Each polyimide shielded bundle was then threaded through a custom 3D printed Microdrive which allows for advancement of each bundle *in vivo* by turning a screw. Each Microdrive was part of a larger custom 3D printed implant (Clear V4 resin, Formlabs, Somerville, MA, USA), with holes for drivable screws (McMaster Carr, Elmhurst, IL, USA) that allowed for advancement of the electrodes before every recording session. Reference and Ground pins were shorted together on each of the EIBs and both EIB grounds were also shorted together. For quality control, channels were then electroplated using a gold solution (Neuralynx, Inc., Bozeman, MT, USA) to match impedance of all channels to 200 kΩ ±100 kΩ. If the implant had >10% (∼6) channels that were either open or shorted, the implant was not used.

#### Surgical Implantation

While maintaining a sterile field, rats were anesthetized using isoflurane (5% for induction, maintained at 1.5-2.5%), placed in a stereotaxic frame, and prepped for Microdrive implantation. Briefly, after shaving and cleaning the skin with betadine, an incision was made to expose the skull. For each hemisphere in mPFC, a hole was drilled for each electrode bundle at + 2.8 AP, ± 0.7 ML. Screws for durability and implant adhesion were also drilled and placed on either side of the orbitofrontal cortex. A stainless-steel screw was fixed to the skull in the cerebellum to act as ground/reference, connected to the EIB ground via an insulated silver wire (A-M Systems cat no. 787000, 0.0130” coated). A thin layer of CCB Metabond was then applied across the skull as well as a small well of dental cement around the implantation site (mPFC). The 64-channel dual bundle/hemisphere implant was then stereotactically lowered to its implant location + 2.8 AP, ± 0.7 ML, and −2.0 DV from brain surface. Dura-Gel (Cambridge NeuroTech) was then applied to the implantation well to seal and stabilize the durotomy. Finally, dental cement was used to fully encase the implant to the skull and rats were then allowed to recover for 1 week before re-training, testing and recording. Just prior to the first recording session, Microdrives were advanced by −1.0 mm to achieve the desired prelimbic coordinates. Waiting to advance until just prior to recording maximizing single unit yields as brain tissue is not yet compromised. For subsequent recording sessions the bundle was only lowered if poor signal was observed, with a maximum goal depth of −1.6 mm.

### Single-unit recordings

During Recording, two Intan RHD 32-Channel Headstages were used, one for each EIB/hemisphere. The Intan RHD Dual Headstage adapter (Part # C3442) was used to connect the two RHD headstages to a single 0.9m SPI cable (#C3203). This then connected to an AlphaComm-I (AlphaOmega) commutator via an Omnetics PZN-12 polarized nano junction. Another SPI cable was then used to interface with an Open Ephys Acquisition Board. The Open Ephys acquisition board also received behavioral events from the Coulbourne operant boxes using TTLs received via an Open Ephys I/O board. All 64 channels of electrophysiological data as well as behavioral TTLs were simultaneously and continuously sampled at 30 kHz using the Open Ephys GUI and raw data was saved via Record Node as a Binary file.

### Spike Sorting

Binary data was then read into MATLAB for offline spike sorting. Single units were sorted using Kilosort3 and then subsequently visually examined using Phy. Single units labeled “Good” or “MUA” by Kilosort were manually inspected in Phy to make sure that the inter-spike-interval (ISI) distribution was Poisson and the unit’s autocorrelation was not violated, and that single unit amplitudes and waveforms did not drift significantly over the course of the recording. Units labelled “MUA” that showed a clear refractory period and non-contaminated waveform were reassigned as “Good”. Spike times from all “Good” single units were then exported to a MATLAB structure for each recording session, containing spike times, and behavioral timestamps.

### Electrode Placement Verification

Once recording was finished electrode placement was verified via electrolytic lesion using a multielectrode NanoZ testing/electroplating device (WhiteMatter, LLC). Rats were anesthetized using isoflurane and then electrolytically lesioned. Rats then received a dose of chloral hydrate (400 mg/kg) and were transcardially perfused with 4% paraformaldehyde. Brains were extracted, post-fixed in 20% sucrose before being sliced in 40 um segments. Brain slices were slide mounted and Nissl stained before being imaged on a Zeiss Apotome microscope to verify correct placements.

### Behavioral Data Analysis

#### Full session accuracy and latency analysis

For each subject we quantified the average accuracy across all trials where there was an optimal choice (low uncertainty blocks). We also quantified action latency (defined as the time to nosepoke following nosepoke illumination) and retrieval latency (defined as the time to enter the feeder trough following nosepoke).

#### Logistic regressions

We employed a logistic regression approach to quantify the extent to which past reinforcement history influenced subject choices (Tsutsui et al., 2017; Grossman et al., 2022).

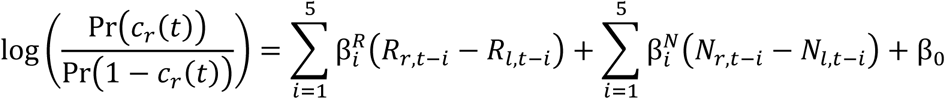

Where c_r_(t) = 1 if right is chosen and 0 if left is chosen on trial (t). Further, R = 1 for rewarded choice and 0 for unrewarded choice, while N = 1 for unrewarded choice and 0 for rewarded choice. Thus, the betas for each of the past five trials capture the difference in likelihood for them to choose the same or different side as a function of reward/no reward by being positive or negative, respectively.

#### Win-stay analysis

We applied a win-stay/lose-shift analysis, split by uncertainty level, to understand what strategies were utilized across blocks. This was done by calculating the probability of staying (subject chose the same side as the prior choice) after a rewarded(win-stay) or unrewarded (lose stay) trial. Note that lose-shift probability here would be 1-lose stay. Like the above analysis, this allows us to assess how recent outcomes influenced choices but in an uncertainty specific manner. Outcome dependent strategy shifts and psilocybin effects were assessed using a two-way repeated measures ANOVA using outcome and session as factors.

#### Choice variability analysis

To quantify choice variability we calculated entropy of choices (Woo et al., 2023) during either high or low uncertainty blocks. This was done using the equation:

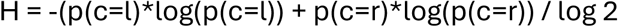

Thus H (entropy) quantifies how unpredictable choice (c) on left (l) and right(r) nosepokes is under the differing contingencies where values of 0 would indicate no entropy (i.e. the animal always picks a particular side) and values of 1 indicate full entropy (i.e. the animal chose both sides equally). Effects of psilocybin were assessed using a paired t-test or Wilcoxon signed rank test depending on normality violations indicated by Shapiro-Wilke tests.

#### Transition analysis

A transition analysis was utilized by assessing choice behavior in the five trials up until a block transition and ten trials after. Given that each block was a minimum of 15 trials these epochs ensure data are not double counted. For transitions from low uncertainty, choices were normalized to the optimal side prior to transition (e.g. c=1 if the right side was chosen when p(R|r) = 0.8 in the pre-transition block).

We used a generalized linear mixed effects model (GLMM) with a binomial link function (lme4 v1.1, lmerTest v3.1 R packages) and compared several models with differing levels of complexity for the fixed effects with likelihood ratio tests (**Statistics Supplement**). This allowed us to test for overfitting and whether additional features such as transition type (low to low vs low to high) or non-linear patterns (e.g. immediate post feedback drops) were important. Our selected model utilized session (baseline vs psilocybin), reversal feedback, and transition trial as fixed effects with rat and reversal id (nested within rat) as random intercepts. This can be written as:

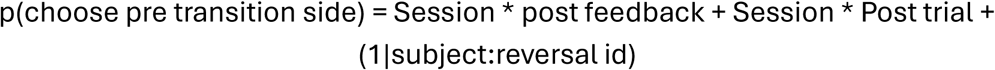

Here post feedback is coded as 0 for trials prior to reversal feedback (i.e. trials −4-0) and 1 for trials after. Post trial is coded to 0 for all trials prior to and including the first feedback trial and increases linearly from 1 thereafter. This parameterization allows estimation of psilocybin effects on asymptotic performance pre-transition (main effect of Session), immediate changes in behavior after transition (Session x post feedback), and the rate of change after transition feedback (Session x post trial), and parallels reversal curve analyses that quantify the asymptotic performance, immediate change after transitions, and post transition choice changes over trials (Findling et al., 2025; Cheng et al., 2025). However the hierarchical framework allowed us to account for the non-independence in the data and unbalanced sampling.

We also took subject averaged 5-trial bins of choice data around transitions (trials −4-0,1-5,6-10). This allowed us to visualize the consistency of the effect across individuals in the asymptotic epoch, and early or middle transition epoch. Psilocybin behavior was compared to baseline using a paired t-test or Wilcoxon signed rank test depending on normality violations indicated by Shapiro-Wilke tests. Given prior work demonstrating acute psilocybin improves rule learning (Torrado-Pacheco et al., 2023), we hypothesized that psilocybin would improve optimal choice behavior at the end of the block (i.e. asymptotic performance). Thus tests with prior directional hypothesis were one-tailed and were two-tailed otherwise.

#### Excluded Data

For latency data, any value greater than 5 standard deviations above the mean was excluded. Due to the order of blocks, one rat only completed 2 trials under high uncertainty during their psilocybin session. This subject was thus excluded from high uncertainty win-stay and choice variability analysis. For transitional analysis, any instances where the full transition window was not present (i.e. < 10 trials post transition) were excluded which resulted in 3 transitions being excluded from psilocybin related data and 4 transitions excluded from saline related data.

### Computational Modeling

#### Model Space

We considered five reinforcement learning (Q-learning) models with varying complexity that were adopted from prior work.

*Model 1-Base*, our simplest model, followed a standard reinforcement Q-learning (RL) framework where action values for each choice V_r_ and Vl are updated using standard Rescorla-Wagner delta learning rule. If the right side was chosen this is expressed as:

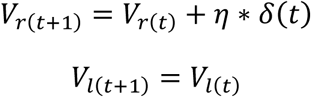

where *δ*(*t*) = *R*(*t*) − *V*_r_(*t*) and R = 0 for unrewarded and R=1 for rewarded trials and η is a free parameter from 0-1 referred to as the learning rate. These value estimates were used as the Q-values to obtain choice probabilities via a logistic function:

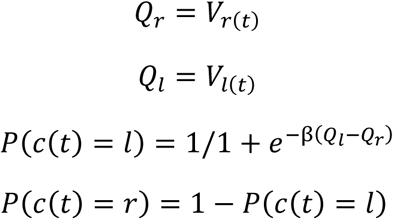

Where β is the inverse temperature which controls the slope of the function.

*Model 2 - Bias* was based off prior work using probabilistic decision-making tasks, which contained an additional bias term in the decision process (Glascher et al., 2009), such that the logistic function is defined as:

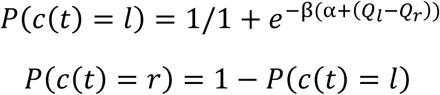

and α is the bias determining whether the function is shifted to a particular side.

*Model 3 – Asymmetric Learning+Bias* utilized the same structure as Model 2 but with a learning rate for each outcome. Thus, the update equation is as follows when the right side is chosen:

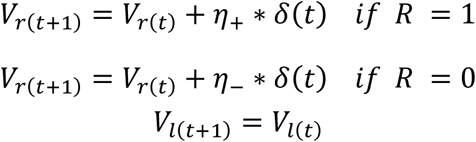

This allows for a different amount of updating to occur as a function of the outcome. This model included bias as in Model 2.

*Model 4 – Split+Bias+Stickiness* utilized the same structure as Model 3 but with a stickiness parameter (κ) to account for perseverative choice. Thus, the following equation is used prior to the logistic function as follows when the right side is chosen:

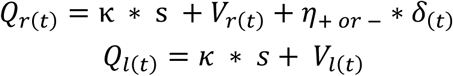

Where s = 1 if Qside = C_t-1_ and 0 otherwise. This allows for the prior choice to increase (or decrease) the likelihood of being repeated in the current choice in an outcome independent manner.

*Model 5 – Meta-learning from uncertainty* This model mirrored recent studies using probabilistic foraging and utilized error-derived expected and unexpected uncertainty estimates to modulate learning (Grossman et al., 2022). This model added additional uncertainty and meta-learning components that modulated updating on a trial by trial basis. Thus, the update equation is as follows when the right side is chosen:

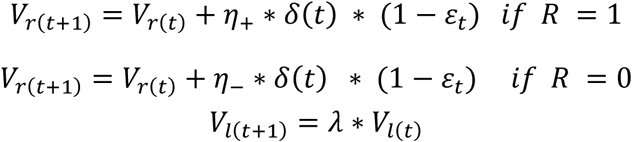

Here, λ is a decay parameter and ε is the expected uncertainty estimate which is derived from unsigned prediction errors as followed:

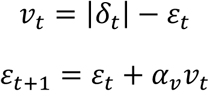

Thus *δ* is the prediction error, *v*_t_ is the estimate of unexpected uncertainty, and α_v_ controls the rate at which the unsigned prediction error is integrated into uncertainty estimates.

Finally, these uncertainty estimates are used to adapt 5_–_:

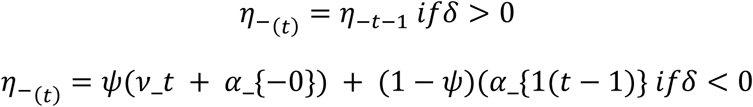

Where α_-0_ is the baseline no reward learning rate and ψ controls the rate at which unexpected uncertainty is integrated into η-after no reward.

The decision is formalized similar to Model 2:

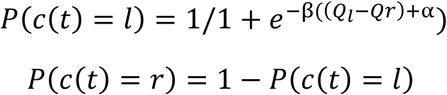

#### Fitting procedures

Models were fit and assessed via Stan (https://mc-stan.org) using the R interface (RStan, version 2.18). Models were constructed hierarchically such that the effect of psilocybin was at the highest level, while subject level effects were below. For each parameter we estimated a population-level mean and standard deviation and subject level parameters were derived from the population level using non-centered parameterization. The treatment effect was modeled as an additive shift from the baseline (e.g. psilocybin learning rate = baseline learning rate + treatment effect), with the baseline condition (‘treatment’) effect constrained to 0. Because bias could be directed towards either side, treatment effects were modeled multiplicatively (psilocybin bias = baseline bias * exp(treatment effect)), effectively modulating the magnitude of bias independent of direction. Thus, the treatment effect estimated reflects the change in each parameter due to psilocybin treatment. This approach accounts for the within-subject nature of the study. Parameters were estimated using weakly informative priors for treatment and subject level parameters that were largely based on other rodent studies (Cheng et al., 2025). Rat level parameters, and group-level means and treatment effects were set as normally distributed with N(0,1) priors. Group level standard deviations were set to Gamma(2,0.5).

Parameter estimates were sampled in unbounded space and later bounded using the fast approximation of the cumulative density function to be between 0-1 (phi_approx) for all parameters except choice bias (α). We also non-centered our parameters to improve sampling. Estimates were generated via Stan through Hamiltonian Markov Chain Monte-Carlo sampling. This was done by running four chains with 2000 samples per chain with the first 1000 samples discarded as burn in. Default configurations were used with the exception of the acceptance rate (0.99) and max tree depth (15) to improve convergence and prevent divergences.

#### Model checks and comparison

Models were checked for convergence using the r-hat values. A criteria of r-hat < 1.1 was used and the maximum observed r-hat was 1.017 for the preferred model. The preferred model was selected based on a combination of pareto-smoothed importance sampling leave one out approach (PSIS-LOO) and its posterior predictive ability to recover psilocybin-related effects (see **Supplemental Data Fig. 2A-B**). While Model 5 produced a similar PSIS-LOO to the winning model, it had more parameters and produced issues with divergences which could not be overcome even with our high acceptance rate or with increasing chain length. This may indicate it was too complex for our given dataset which was considerably smaller than the dataset this model was based off due to the nature of our acute dosing study.

#### Posterior checks and parameter estimates

We assessed whether our models could recapitulate key trends in our data, particularly around transitions. To do this we simulated an agent on the task using the same trial structure a given animal received using the mean of that given animal’s posterior estimates for each variable. This was done 4000 times per agent and we then took the choice probability average across all runs to create a simulated data set that matched the exact number of trials from the observed data.

For parameters estimates we assessed the 4000 posterior draws for the group level effect of psilocybin. To assess group differences, we used ArViz (version 0.21) to calculate the 80/90/95% highest density intervals (HDI) of these distributions. A significant change from treatment was determined if the 80% HDI did not include a value of 0 (Hervig et al., 2024). We also quantified the strength of evidence for a shift in parameter values using directional bayes factor (dBF; Kass C Rafferty, 1995; Cheng et al., 2025).

### Neuronal data analysis

#### Population level neuronal activity

To assess how outcome related neural activity was modulated by treatment, we calculated peri-event averages in spike rate across all neurons around the 1 sec before and 1.5-sec after action execution (Action-Outcome). To do this we quantified firing rates of units by binning spike counts using a sliding time bin (200 msec bin width, 50 msec bin step). To assess outcome related shifts in population activity, z-scored bin counts were averaged over reward or no reward trials. For neural analyses we found that neurons with high firing rates (>20 Hz; Park & Moghaddam 2017) made up only approximately 6% of the recorded units. This was also in agreement with other approaches (Le Merre et al., 2026) such as subclassification of units as wide width (ww) or narrow width (nw) based on peak to trough durations (5-6% classified as nw using a 0.43 ms threshold; **Supplemental Data Fig. 3B**).

Given we were interested in general PFC neural activity we opted to include these units in all analyses.

To assess how session (treatment) altered the trajectory of population unit activity across the epoch we modeled mean firing rates using a mixed effects model (R lme4 package, version 1.1). This approach allowed us to additionally account for random effects between individual rats and units. Thus the model was defined as follows:

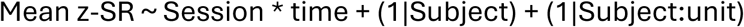

Where SR is the mean z-normalized spike rate, Session was either baseline or psilocybin and time was the time bin. Because we noted non-linear activity fluctuations around action/outcome, we modeled time as linear or as a 2^nd^ or 3^rd^ order polynomial (e.g. poly(time,3)). These models were compared using likelihood-ratio tests to validate model improvements from the added polynomial complexity. Subject and unit are random effects which allow for a random intercept for each unit and animal. P values for interactions and main effects were obtained via the lmerTest package (version 3.1).

### Hierarchical clustering of outcome related activity

To examine the variability in response to outcome we concatenated the mean z-normalized peri-action activity for reward and no reward trials for each unit over the post outcome period (0.05-1.5 s). This created a matrix where each unit was a single vector of reward and no reward response. We then computed Pearson correlations (*r*) between pairs of units to create a correlation matrix. To convert the correlation values matrix into a distance matrix for hierarchical clustering we transformed correlation values such that distance = 1-*r*.

Unsupervised agglomerative hierarchical clustering was performed using complete linkage on distance profiles for each unit (scipy linkage). Prior work in human and rodent cortex has indicated the number of unit clusters typically ranges around 4-6 (Aponik-Gremillion et al., 2022; Le Merre et al., 2026). We utilized this knowledge and the Thorndike method to determine how many clusters were present (Thorndike 1953; Le Merre et al., 2026). The majority of changes in distance were confined to the first five clusters in the dendrogram (68% of maximum distance; **Supplemental Data Fig. 4B-C**).

To assess the robustness of our clustering solution, we used a leave-one-out procedure (Aponik-Gremillion et al., 2022). For each unit, we removed said unit, recomputed the clustering procedure on the remaining units, and compared the resulting cluster groupings to those seen in the full dataset using the Adjusted Rand Index (Hubert & Aribie, 1985; ARI). ARI values range from −0.5-1, with −0.5 indicating below chance, 0 indicating chance and 1 indicating identical clustering to the original clustering solution.

### Characterization and comparison of functional clusters

After cluster assignment, we characterized the functional properties by examining the average response profile for rewarded and unrewarded outcomes. Cluster were categorized by their dominant response patterns, specifically whether they increased or decreased activity post-outcome, and whether this response was elevated for reward versus no reward. Thus clusters which produced elevated firing rates for reward over no reward were classified as reward preferring, clusters which produced elevated firing rates for no reward over reward were classified as no reward preferring, and clusters which showed ramping activity that decayed at outcome were classified as action-decay.

To test if psilocybin altered the relative prevalence of different functional subclusters, we compared the composition of clusters between sessions by calculating the number of units assigned to each cluster and normalizing to the total number of units recorded for that session. Statistical significance was assessed using Fisher’s exact tests for each cluster with a Benjamin-Hochberg correction applied to p-values. To test if psilocybin altered relative preference magnitude, we calculated the difference in reward and no reward average z-score for 0.5 second chunks after action. These values were compared via mixed effects models with fixed effects of session and bin and unit nested within subject as random effects. Significant effects were followed by post-hoc t-tests with Benjamin-Hochberg corrections if needed.

### Regression models of neuronal data

We fit regression models to assess how both task variables and decision values (Donahue, Seo, & Lee, 2013; Shin et al., 2021) were encoded by unit spike activity in the mPFC. We fit two regressions designed to predict the spike rate of an individual unit as a function of either task features (choices and outcomes) or value signals derived from reinforcement learning modeling (see below). Given that psilocybin decreased the number of trials completed (Baseline Mean ± SEM= 166±18.2, Psilocybin Mean ± SEM= 99±18.7; p < .001, paired t-test), and to match other studies comparing drug to non-drug encoding (Cheng et al., 2025), we matched the number of trials for a subjects’ baseline session to the corresponding subject’s number of trials in their psilocybin session. Furthermore, given that having fewer trials could also enhance trial related correlations, we took a more conservative approach by first regressing out the effect of trial number on spike count and running future encoding on the residuals. Models were fit using python’s statsmodels package (v0.14).

As with population analyses, we quantified firing rates of units by binning spike counts using a sliding time bin (200 msec bin width, 50 msec bin step) during the Action-Outcome epoch. Models were fit for each bin around the windows of interest across trials (t). We adopted the same method as others (Shin et al., 2021) to control autocorrelation among spikes by including the spike counts for three prior trials as autoregressive components in the model. Thus for each time bin within the window the task variable model was defined as:

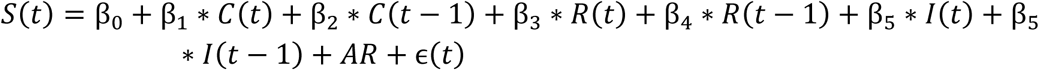

Where S(t) is the residual spike counts for a given bin, *β*_1_ through *β*_5_ are coefficients for choices (C), outcomes (O), or the interaction between current or prior choice and outcome (I). AR is defined as the following and treated as a continuous variable:

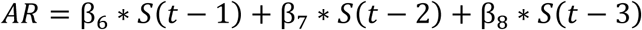

To assess how value-related signals were encoded we used the following choice-value model:

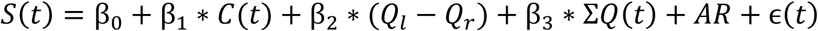

Here, the C(t) is the chosen direction, utilized to account for the fact that value signals can be correlated with choice. Ql-Q_r_ is the difference in value between the two options, and ΣQ(t) is the total value for both choices on each trial. Both Q-related values were z normalized for each subject and treated as continuous variables.

After fitting the model, t-statistics were used to determine significance of betas by comparing the beta to a null of 0. Corresponding p values for each of the predictors were obtained. We used a permutation-based approach to determine a multiple comparison adjusted threshold for encoding (Tsutsui et al., 2016; Costa et al., 2019). To do this we refit models for each neuron 250 times for a given factor after randomly shuffling the factor of interest each time. We saved the largest number of consecutive time bins that achieved significance for each factor and used these permutations across units to generate a null distribution. We then used this null distribution to determine the value which defined the 95% percentile, this resulted in a value of 4-5 across factors which is in line with other studies using similar bin widths and step sizes (Costa et al., 2019; Tang et al., 2022). Thus we opted to use a consecutive threshold of 5 contiguous bins (i.e. unit was considered to encode a given factor if it maintained >=5 consecutive p values of less than .05).

To assess encoding over the peri-action epoch across encoders, we took the task (or value) responsive neurons and calculated the percent of this population which showed encoding and compared to chance levels at each time bin using a binomial test (Costa et al., 2019). Binomial test p-values (compared to 0.05) were post-hoc corrected using the Benjamin-Hochburg method and significance was determined at a p<.01 significance level.

### Comparisons of psilocybin effects on encoding

We used two complementary approaches to assess if psilocybin altered the time-course of a factor encoding compared to baseline: cluster corrected chi-squared tests and mixed effects models. Chi-squared tests were applied at each time bin, providing fine-grained temporal resolution. Mixed-effects models were applied to each units average encoding across broader 0.5 s epochs, which gave greater statistical power to detect sustained shifts in encoding magnitude. Together these approaches provide both the temporal specificity to localize when encoding differences emerge and the sensitivity to characterize more diffuse changes in overall magnitude.

Chi-squared tests were performed over all time-bins within the peri-action period (−1-1.5 s). To control for multiple comparisons we employed a cluster-based permutation approach. Session labels were randomly shuffled and chi-squared tests were re-run across all time-bins. The longest contiguous run of bins with p < 0.05 was recorded. This procedure was repeated 1000 times to generate a null distribution of maximum cluster lengths. The cluster threshold was defined as the minimum cluster length that occurred in fewer than 5% of the shuffle iterations. Observed clusters meeting or exceeding this threshold were considered significant.

Mixed effects models (R lme4 package, version 1.1) assessed how treatment effects differed over time. This approach also allowed us to account for random effects between individual rats and units (Cheng et al., 2025). Thus the model was defined as follows:

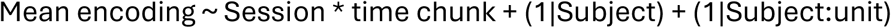

The fixed effects here are Session (psilocybin versus baseline) and time chunk (relative time around event) and the related interactions. Subject and unit are random effects which allow for a random intercept for each unit and animal. In the case of significant beta(s) for the interaction of time with session, we performed post-hoc Session contrasts for each time chunk via the emmeans package (version 1.11) using the Benjamin-Hochberg correction.

### Decoding analysis

#### Preprocessing

Binned spike data (200 ms bin, 50 ms step) was first z-normalized for each unit. To isolate neural activity related to the variable of interest independent of task structure and history, we regressed out confounding covariates such as trial, block type, and reward or choice. For example if we were decoding choice, for each unit at each timepoint we fit a linear regression model to predict spike rate with trial, block type, and reward as covariates and saved the residuals from this regression. We then ran decoding analyses on these residuals.

#### Pseudosession construction and decoding procedure

We used a pseudo-session approach to decode either choice direction or trial outcome from mPFC population activity across the peri action epoch. At each time bin we randomly sampled, without replacement, 10 trials each for each level of the factor we were decoding (e.g. 10 left choices, 10 right choices) for each unit. For each sampled pseudo-session, we used the corresponding activity of each unit at the given peri-action time bin for those trials. Thus this created a feature matrix of 20 trials by n-units used for decoding (see below) where the target was choice or outcome and features were z-normalized spike counts for each unit in a given time bin.

Decoding analysis was done by training a L2-regularized (C=1) logistic regression classifier (scikitlearn version 1.5.1) to predict choice or outcome. L2 regularization was used to prevent overfitting given that the number of features could substantially exceed the number of pseudotrials at certain population sizes. We then used leave-one-out cross validation by training the classifier on N-1 pseudotrials and testing on the held out trial. Classification accuracy was computed from the mean accuracy on the held out trial across all folds. We repeated these procedures 20 times per condition to attain stable estimates of decoding accuracy, with final accuracy reported by the mean and confidence interval of the average per each run (i.e. n=20). To determine chance levels we ran decoding analysis in the same way as above but permuted the target labels prior to training.

#### Population size analysis

We systematically varied the number of units used as features in the decoding. This was done by randomly sampling populations of 10, 50, 100, 150, 200, and all available units (full), and performing the decoding procedures for each population size. This allowed us to determine the relationship between population size and decoding ability and whether psilocybin shifted this pattern.

#### Subcluster ablation tests

To assess which neuronal subclusters contributed the most to decoding accuracy we performed an ablation analysis based on the functional subclusters outlined in hierarchical clustering analysis. For each subcluster we repeated the full decoding analysis after removing neurons belonging to that subcluster. The contribution of each subcluster to decoding was quantified by comparing the accuracy of the decoder without these units to that of the full population Δ %Accuracy = (Ablated accuracy – Full Population Accuracy) * 100. Because the sizes of subcluster could differ between clusters and sessions, and this could effect changes in accuracy, we normalized this change by dividing it by the number of units in the subcluster, effectively capturing the change in accuracy per unit lost.

#### Statistical Testing

We compared decoding accuracy for either the pre-action (−1-0 s) or post action (0.05-1.5 s) period for population size and ablation analyses. Statistical significance was determined by comparison to permutation (shuffle) related performance (to assess if decoding was above chance) or comparing baseline to psilocybin to assess effects of treatment. This was done using a two-way ANOVA with effects of either session (or shuffle) and population size (or subcluster) with post hoc contrasts using Benjamin-Hochberg correction where required.

### Data Visualization

All data were plotted using Python or R packages/libraries of Seaborn (v 0.13, Waskom 2021), Upset (v 0.9, Lex, 2014), Matplotlib (v 3.9, Hunter et al., 2007), and ggplot (v 3.5, Wickham, 2016).

## Supporting information

Statistics Tables

## Acknowledgements

The authors that Dr. Merridee J. Lefner for their thoughtful comments on the manuscript.

## Ethics Declarations

### Contributions

Conceptualization and design: A.T.P and B.M. Data collection: A.T.P and R.O. Data Analysis: D.S.J., A.T.P. and A.J.L. Funding Acquisition: B.M. Supervision: B.M. and A.J.L. Wrote the first version of the manuscript: D.S.J and B.M. Edited the manuscript: D.S.J, A.T.P., R.O., A.J.L, B.M.

