## Supplemental Figures for "Psilocybin asymmetrically modulates outcome-based choice and cortical processing under uncertainty"

**Supplemental Data.**

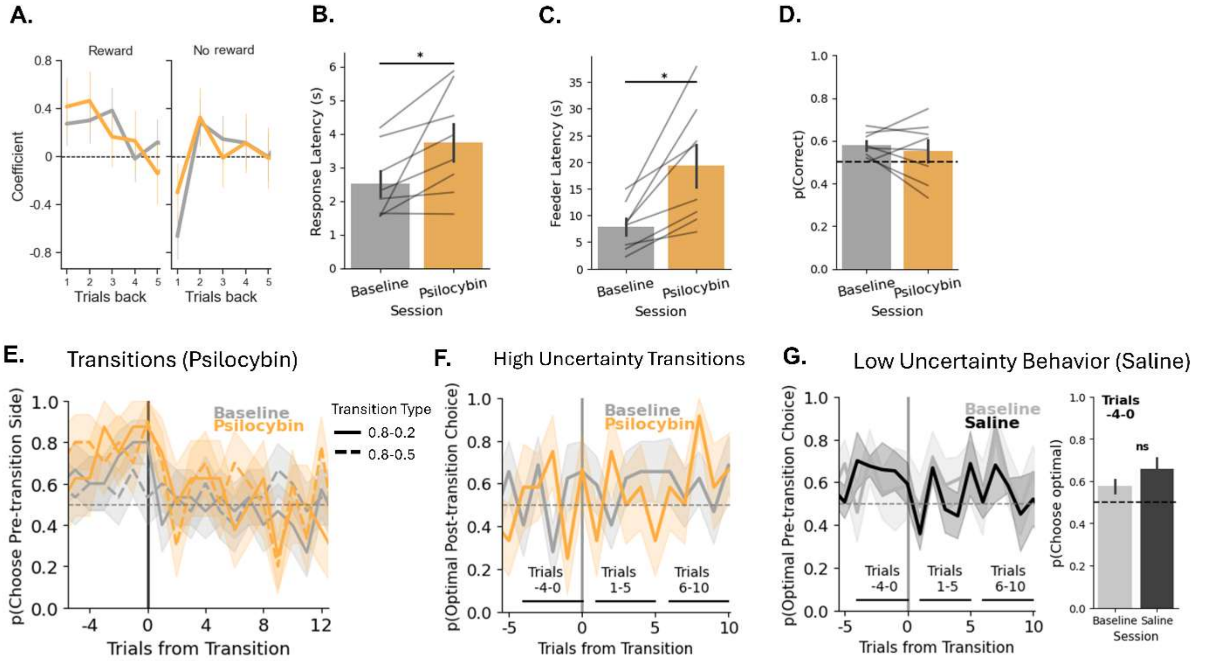


**Supplemental Data 1.** Supplemental behavioral and saline related data. **A.** Coefficients for logistic regressions of choice following reward and no reward for baseline and psilocybin sessions. Data reflect mean and 95% confidence intervals. **B.** Latency to execute actions for baseline and psilocybin sessions. Psilocybin increased latency to execute a response (p=.031, paired t-test). Lines indicate individual rats (n=8). **C.** Latency to enter the feeder following actions for baseline and psilocybin sessions. Psilocybin increased latency (p=.006, paired t-test). Lines indicate individual rats (n=8). **D.** Average accuracy across blocks for session, no significant differences were observed (p=.54, paired t-test). Lines indicate individual rats (n=8). **E.** Transition analysis for low uncertainty transitions split by transition subtype (low-low, low-high uncertainty). Transition type was not a significant indicator of behavior within the (-4-10 trial time window, see **Statistics Supplement**). **F.** Transitions from high uncertainty indicate chance choice levels pre-transition and non-significant changes in the initial 10 trials into low uncertainty (n=7 rats). Note because trials -4-0 have no 'optimal' option, all choices are compared to optimal option post-transition. **G.** Similar behavioral patterns between saline and the corresponding baseline (pre-saline) session were observed for transition analysis from low uncertainty. N=4-5 rats. **Right-** Optimal pre-transition performance (trials -4-0) was not influenced by saline (p=.31, paired t-test).

**A.**

| | Model | Params | PSIS-LOOIC (Psilocybin session only) | PSIS-LOOIC (Both Session) |
|---|---|---|---|---|
| 1 | Base | β, η | 1043 | 2768 |
| 2 | Bias | α, β, η | 1010 | 2676 |
| **3** | **Bias+ Asymmetric learning** | **α, β, η+, η-** | **989.7** | **2647** |
| 4 | Sticky | α, β, η+, η-, κ | 1005 | 2562 |
| 5 | Uncertainty-based meta learning (74 divergences) | α, β, η+, η-, $\eta_0$, ω, λ | 987.4 | 2619 |

**B.**

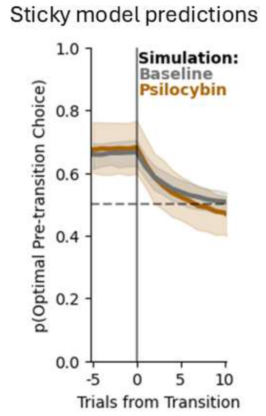


**Supplemental Data 2.** Computational models of behavior, pareto smoothed importance sampling values (PSIS-LOO), and model predictions from the 'sticky' model. **A.** Outline of models tested, free parameters, and PSIS-LOO for the whole dataset and psilocybin specific data. Bolded model was our chosen model. Only Model 5 produced divergence issues and was considerably more complex than our chosen model. **B.** Prediction from the 'sticky' model. Although the PSIS-LOO was lower for the full dataset it performed poorly at recovering the psilocybin data around transitions.

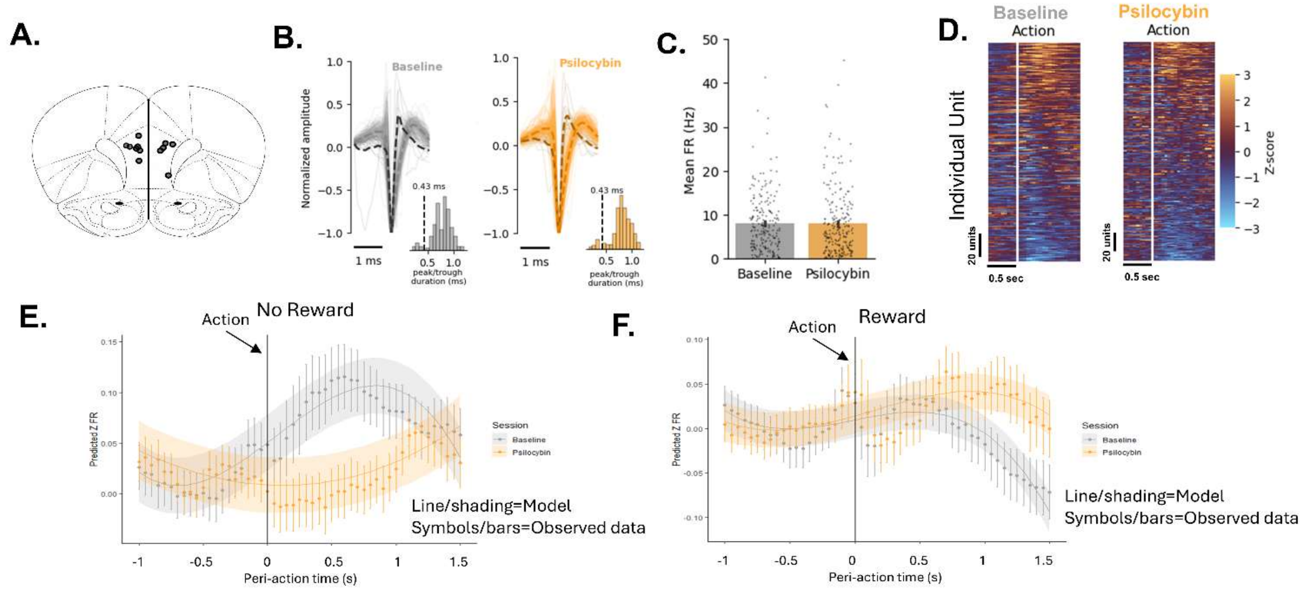


**Supplemental Data 3.** Supplemental data related to single units and population activity. **A.** Hitmap of electrode placements. **B.** Mean normalized waveforms for sorted units in baseline and psilocybin sessions. Color shading indicates whether unit was narrow width (nw; darker) or wide width (ww; lighter) based on a peak to trough duration. Solid lines are individual units traces while dashed lines are the mean for the nw or ww subsets. Bottom histograms show the distribution of peak to trough durations for all units and the threshold for nw/ww determination at 0.43 ms (n=207 units for baseline and n=222 units for psilocybin from 7-8 rats). **C.** Mean basal firing rate of units in baseline and psilocybin sessions (p=.84, unpaired t-test). **D.** Mean z-score for peri-action activity across units. Rows are individual unit z-normalized spike rate and columns are time bins. White vertical line indicates action execution. **E-F.** Overlay of model predictions from a 3$^{rd}$ order polynomial mixed effects model (shading) onto unit averages (circles) for unrewarded and rewarded trials (mean ± SEM).

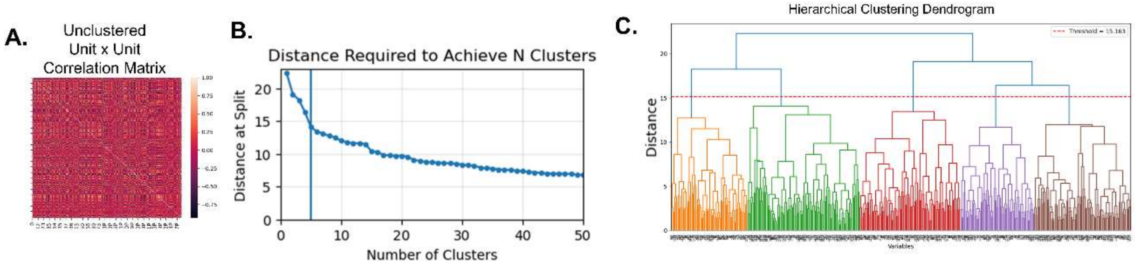


**Supplemental Data 4.** Supplementary data for clustering analysis. **A.** Unit by unit correlation matrix prior to sorting units based on hierarchical clustering. **B.** Distance change for each cluster split for 1-50 clustering solutions. The ‘elbow’ point appears at about 5-6 clusters. Blue line indicates our chosen cluster number (5). **C.** Dendrogram of clusters shows the height used to create our 5 groups as well as their hierarchical representation.

‘

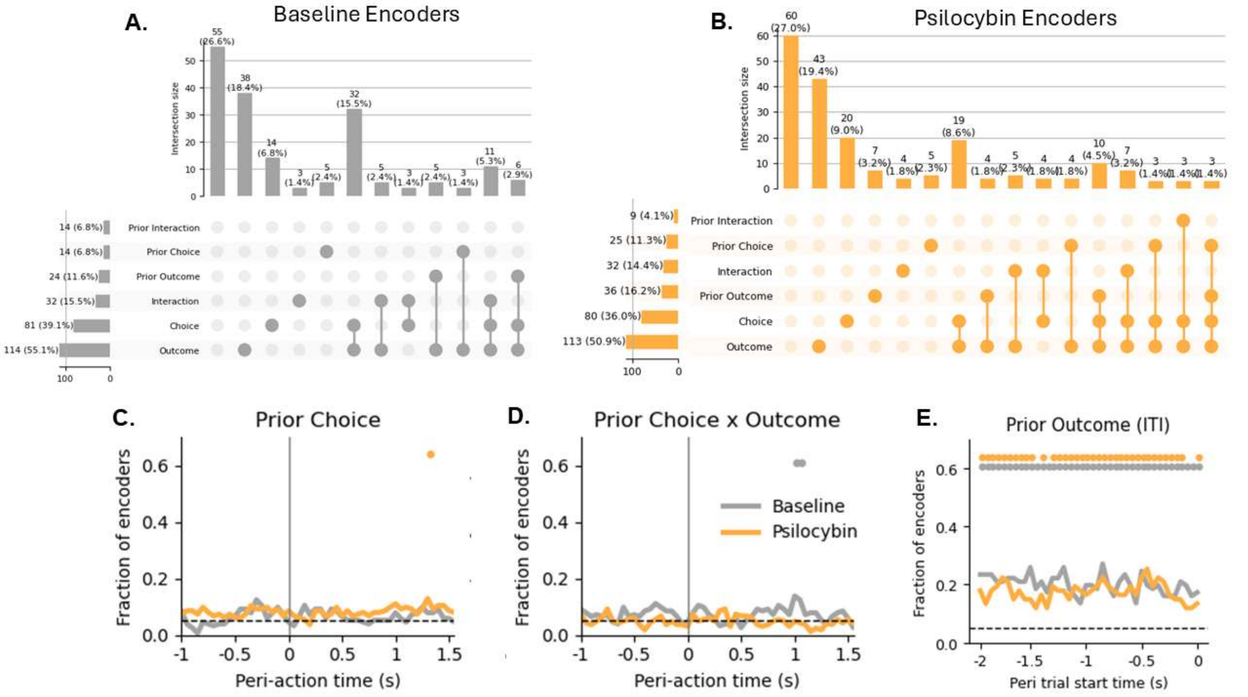


**Supplemental Data 5.** Supplementary data for task encoding models. **A.** Upset plot which illustrates the percentages and selectiveness of unit encoding across outcome and choice related encoding for Baseline sessions. Filled circles indicate which feature(s) units encoded and correspond to the bar above. Horizontal bar plots on the side indicate percentages which encoded a given feature either alone or in combination with other features. To simplify plot, only variable combinations with >=3 encoders were included. **B.** Upset plot which illustrates the percentages and selectiveness of unit encoding across outcome and choice related encoding for Psilocybin sessions. Filled circles indicate which feature(s) units encoded and correspond to the bar above. Horizontal bar plots on the side indicate percentages which encoded a given feature either alone or in combination with other features. To simplify plot, only variable combinations with >=3 encoders were included. **C.** Fraction of task encoders showing significant prior choice encoding at each time bin over the peri-action period. **D.** Fraction of task encoders showing significant prior choice by prior outcome encoding at each time bin over the peri-action period. **E.** Fraction of task encoders showing significant prior outcome encoding at each time bin in the last 2 seconds of the ITI. Colored circles above plots signify significant binomial test ($p<.01$ Benjamin-Hochberg corrected) for baseline or psilocybin.

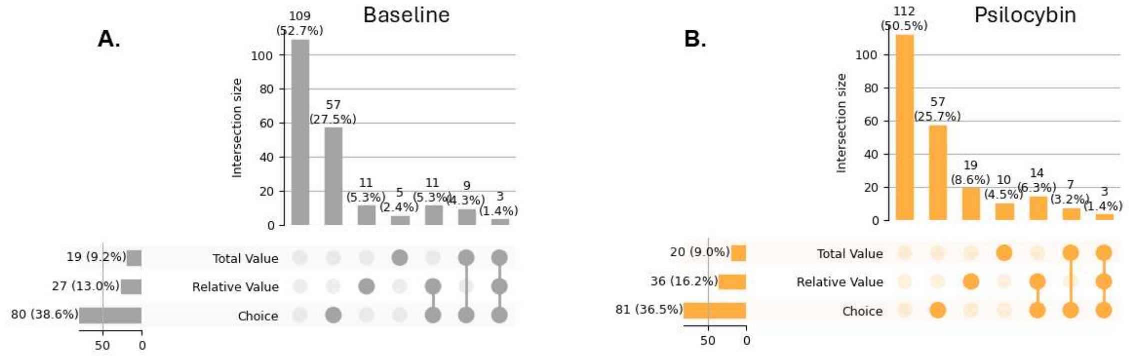


**Supplemental Data 6.** Supplementary data for choice-value encoding models. **A.** Upset plot which illustrates the percentages and selectiveness of unit encoding across choice, relative value, or total value related encoding for Baseline sessions. Filled circles indicate which feature units encoded and correspond to the bar above. Horizontal bar plots on the side indicate percentages which encoded a given feature either alone or in combination with other features. To simplify plot, only variable combinations with >=3 encoders were included. **B.** Upset plot which illustrates the percentages and selectiveness of unit encoding across choice, relative value, or total value related encoding for Psilocybin sessions. Filled circles indicate which feature units encoded and correspond to the bar above. Horizontal bar plots on the side indicate percentages which encoded a given feature either alone or in combination with other features. To simplify plot, only variable combinations with >=3 encoders were included.

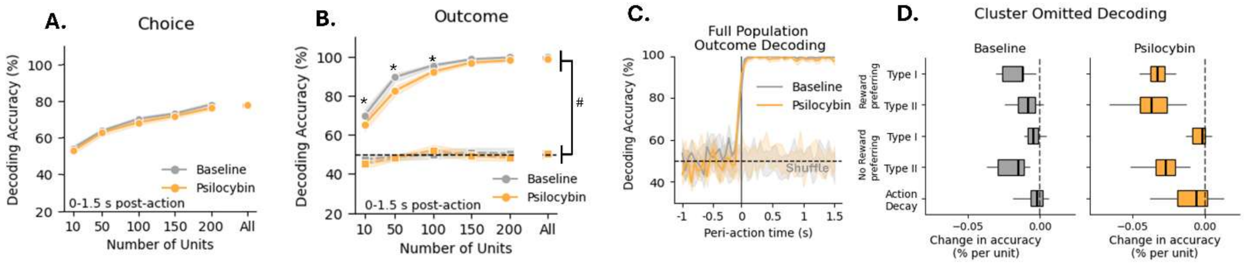

**Supplemental Data 7.** Supplementary data for decoding analyses. **A.** Decoding accuracy for choice across pseudopopulations during the 1.5 second epoch *after* action execution. No significant differences were observed (ANOVA session or session x size p values >.13). **B.** Decoding outcome after action execution was not affected by psilocybin at larger pseudopopulation sizes (n>100 units; mean ± 95%CI, *p<.05 corrected effect of psilocybin at given population size). Squares near the 50% chance line are the results from shuffle analysis ($^{\#}$p<.05 versus shuffle). **C.** Full population decoding (mean ± 95%CI, n = 20 runs with 207-222 units) demonstrated that decoding performance for outcome was highest after action (at outcome delivery) for both sessions. Lighter shading indicates mean ± 95%CI for shuffle runs. **D.** Box plots (median ± whiskers for the 5-95 percentile) showing effects of removing specific ensembles from outcome decoding analysis. Action decay neurons generally had no difference or less effect on decoding accuracy than other clusters (**Statistics Supplement**). The largest losses in accuracy were observed when rewarded or unrewarded preferring neurons were omitted.
