## Supplementary material for "Psilocybin asymmetrically modulates outcome-based choice and cortical processing under uncertainty": Statistics Tables

**Figure 1**

| Panel | Stats | Post-hoc tests/ P Adjustments | Notes |
|---|---|---|---|
| 1D | statsmodels RM ANOVA F DFn DFd p<br>Session 0.3373 1.0000 7.0000 0.5796<br>Outcome 23.2397 1.0000 7.0000 0.0019<br>Session:Outcome 0.2089 1.0000 7.0000 0.6615 | N/A (only 2 levels) | |
| 1E | statsmodels RM ANOVA F DFn DFd p<br>Session 0.0188 1.0000 6.0000 0.8954<br>Outcome 41.0255 1.0000 6.0000 0.0007<br>Session:Outcome 2.0467 1.0000 6.0000 0.2025 | N/A (only 2 levels) | 1 rat excluded (only 2 high uncertainty trials) |
| 1F | scipy paired t<br>t(7) -0.65 p = 0.53 | NA | |
| 1G | scipy paired t<br>t(6)=-3.22 p = 0.018 | NA | 1 rat excluded (only 2 high uncertainty trials) |
| 1H | Model Selection for 1H<br>Data: dffilt<br>Models:<br>m1: value ~ post_trial * Session + (1 \| rat:rev_id)<br>m3: value ~ Session * phase + Session * post0 + (1 \| rat:rev_id)<br>m2: value ~ cert_switch * post_trial * Session + (1 \| rat:rev_id)<br>mfull: value ~ Session * phase + Session * post0 + post0 * cert_switch + (1 \| rat:rev_id)<br>npar AIC BIC logLik -2*log(L) Chisq Df Pr(>Chisq)<br>m1 5 951.07 973.97 -470.54 941.07<br>m3 7 948.01 980.06 -467.00 934.01 7.0638 2 0.02925 *<br>m2 9 955.24 996.45 -468.62 937.24 0.0000 2 1.00000<br>mfull 9 948.78 989.99 -465.39 930.78 6.4601 0<br><br>selected model<br>Generalized linear mixed model fit by maximum likelihood (Laplace Approximation)<br>Estimate Std. Error z value Pr(>\|z\|)<br>(Intercept) 6.500e-01 2.064e-01 3.150 0.00163<br>SessionPsilocybin 7.480e-01 3.617e-01 2.068 0.03868<br>phase -6.257e-01 2.857e-01 -2.190 0.02850<br>post0 2.953e-08 4.182e-02 0.000 1.00000<br>SessionPsilocybin:phase -2.183e-01 4.864e-01 -0.449 0.65351<br>SessionPsilocybin:post0 -8.612e-02 6.865e-02 -1.254 0.20969 | N/A (only 2 levels) | Liklihood ratio test for model selection<br><br>Models fit on raw choice data so binomial (logit) family was used<br><br>post trial coded as n trials after transition<br>Session coded as baseline or psilocybin<br>phase coded as 1 if post transition feedback has occurred<br>(allows an immediate 'drop' follow feedback)<br>post0 coded as trial n after first reversal feedback<br>(linear change after accounting for initial drop) |
| 1I | scipy<br>paired t t(7) = -2.44 p = 0.022 | NA | one tailed |
| 1J | scipy<br>paired t t(7) = -2.57 p = 0.037 | NA | |
| 1K | scipy<br>paired t t(7) = -0.39 p = 0.7 | NA | |

**Figure 2**

<table>
<tr><th>Panel</th><th>Stats</th><th>Notes</th></tr>
<tr><td>2C</td><td>Linear mixed model fit by REML. t-tests use Satterthwaite's method<br>Estimate Std. Error df t value Pr(>|t|)<br>(Intercept) 0.653221 0.024043 10.289465 27.169 6.45e-11<br>SessionPsilocybin 0.095299 0.015616 227.000001 6.103 4.46e-09<br>phase -0.062709 0.018236 227.000001 -3.439 0.000695<br>post0 -0.011394 0.002718 227.000001 -4.191 3.97e-05<br>SessionPsilocybin:phase -0.025163 0.025789 227.000001 -0.976 0.330230<br>SessionPsilocybin:post0 -0.011837 0.003844 227.000001 -3.079 0.002333</td><td>Using lme4 with gaussian family<br>Model structure was value ~ Session * phase + session *post0<br><br>we drop the reversal id from the random effects struture since we are taking averages of agents here.</td></tr>
<tr><td>2d</td><td>paired t t = -2.36 p = 0.025</td><td>using ttest_rel in scipy<br>one tailed</td></tr>
<tr><td>2e</td><td>paired t t =-3.74 p = 0.004</td><td>using ttest_rel in scipy<br>one tailed</td></tr>
</table>

**Figure 3**

| Panel | Stats | Post-hoc tests/ P Adjustments | Notes |
|---|---|---|---|
| 3A | Model Selection:<br>model: value ~ variable * as.factor(Session) + (1 \| id) + (1 \| id:unit)<br>model2: value ~ poly(variable, 2) * as.factor(Session) + (1 \| id) + (1 \| id:unit)<br>model3: value ~ poly(variable, 3) * as.factor(Session) + (1 \| id) + (1 \| id:unit)<br>npar AIC BIC logLik -2*log(L) Chisq Df Pr(>Chisq)<br>model 7 18329 18385 -9157.5 18315<br>model2 9 18309 18381 -9145.7 18291 23.529 2 7.775e-06 ***<br>model3 11 18294 18382 -9135.8 18272 19.700 2 5.275e-05 ***<br><br>Linear mixed model fit by REML. t-tests use Satterthwaite's method<br>Fixed effects:<br>Estimate Std. Error df t value Pr(>\|t\|)<br>(Intercept) -2.151e-03 1.918e-02 5.959e+00 -0.112 0.914370<br>poly(variable, 3)1 -2.257e+00 5.171e-01 2.144e+04 -4.365 1.28e-05<br>poly(variable, 3)2 -2.427e+00 5.171e-01 2.144e+04 -4.694 2.70e-06<br>poly(variable, 3)3 -1.954e+00 5.171e-01 2.144e+04 -3.778 0.000158<br>as.factor(Session)Psilocybin 2.235e-02 1.533e-02 4.235e+02 1.458 0.145684<br>poly(variable, 3)1:as.factor(Session)Psilocybin 4.175e+00 7.189e-01 2.144e+04 5.808 6.43e-09<br>poly(variable, 3)2:as.factor(Session)Psilocybin 1.810e+00 7.189e-01 2.144e+04 2.518 0.011819<br>poly(variable, 3)3:as.factor(Session)Psilocybin 7.904e-01 7.189e-01 2.144e+04 1.099 0.271588 | | id=subject<br>variable=timebin |
| 3B | Model Selection:<br>model: value ~ variable * as.factor(Session) + (1 \| id) + (1 \| id:unit)<br>model2: value ~ poly(variable, 2) * as.factor(Session) + (1 \| id) + (1 \| id:unit)<br>model3: value ~ poly(variable, 3) * as.factor(Session) + (1 \| id) + (1 \| id:unit)<br>npar AIC BIC logLik -2*log(L) Chisq Df Pr(>Chisq)<br>model 7 14282 14338 -7134.0 14268<br>model2 9 14244 14316 -7112.9 14226 42.098 2 7.219e-10 ***<br>model3 11 14218 14306 -7098.3 14196 29.328 2 4.281e-07 ***<br><br>Linear mixed model fit by REML. t-tests use Satterthwaite's method<br>Estimate Std. Error df t value Pr(>\|t\|)<br>(Intercept) 5.837e-02 2.713e-02 6.436e+00 2.152 0.0719<br>poly(variable, 3)1 4.121e+00 4.694e-01 2.144e+04 8.778 < 2e-16<br>poly(variable, 3)2 -2.106e+00 4.694e-01 2.144e+04 -4.487 7.26e-06<br>poly(variable, 3)3 -2.542e+00 4.694e-01 2.144e+04 -5.414 6.22e-08<br>as.factor(Session)Psilocybin -3.360e-02 1.671e-02 4.225e+02 -2.010 0.0450<br>poly(variable, 3)1:as.factor(Session)Psilocybin -3.144e+00 6.526e-01 2.144e+04 -4.817 1.47e-06<br>poly(variable, 3)2:as.factor(Session)Psilocybin 4.235e+00 6.526e-01 2.144e+04 6.490 8.78e-11<br>poly(variable, 3)3:as.factor(Session)Psilocybin 2.612e+00 6.526e-01 2.144e+04 4.003 6.28e-05 | | id=subject<br>variable=timebin<br><br>used bobyqa optimizer<br>to improve convergence |
| 3D | Fisher Exact test (scipy)<br>Type I Cluster: OR = 0.49, p=0.0024<br>Type II Cluster: OR = 0.86, p=0.6041 | Type I BH adjusted p = .011<br>Type II BH adjusted p = .60 | |
| 3F | Type I: Mixed Effects (lmer +lmerTest)<br>Type III Analysis of Variance Table with Satterthwaite's method<br>Sum Sq Mean Sq NumDF DenDF F value Pr(>F)<br>as.factor(chunk) 9.7179 4.8590 2 206 51.0196 <2e-16<br>as.factor(Session) 0.0217 0.0217 1 103 0.2282 0.6339<br>as.factor(chunk):as.factor(Session) 0.0417 0.0208 2 206 0.2187 0.8037<br><br>Type III Analysis of Variance Table with Satterthwaite's method<br>Type II: Mixed Effects (lmer +lmerTest)<br>Sum Sq Mean Sq NumDF DenDF F value Pr(>F)<br>as.factor(chunk) 11.1854 5.5927 2 138 37.3100 1.114e-13<br>as.factor(Session) 0.0783 0.0783 1 69 0.5226 0.4722<br>as.factor(chunk):as.factor(Session) 0.4667 0.2334 2 138 1.5567 0.2145 | BH FDR Corrected p for emmeans contrasts:<br>0.5-0.1 and 1-1.5 vs. 0.0-0.5 p <.001<br>0.5-1 vs. 1-1.5 p = .88<br><br>BH FDR Corrected p for emmeans contrasts:<br>0.5-0.1 and 1-1.5 vs. 0.0-0.5 p <.001<br>0.5-1 vs. 1-1.5 p = <.001 | chunk=time bin<br>0-0.5<br>0.5-1<br>1-1.5 |
| 3G | Fisher Exact test (scipy)<br>Type I Cluster: OR = 1.8, p=0.0024<br>Type II Cluster: OR = 0.82, p=0.48 | Type I BH adjusted p = .036<br>Type II BH adjusted p = .59 | |
| 3I | Type I: Mixed Effects (lmer +lmerTest)<br>Type III Analysis of Variance Table with Satterthwaite's method<br>Sum Sq Mean Sq NumDF DenDF F value Pr(>F)<br>as.factor(chunk) 2.19562 1.09781 2 184 12.6005 7.437e-06<br>as.factor(Session) 0.11325 0.11325 1 92 1.2999 0.2572<br>as.factor(chunk):as.factor(Session) 0.27387 0.13693 2 184 1.5717 0.2105<br><br>Type II: Mixed Effects (lmer +lmerTest)<br>Type III Analysis of Variance Table with Satterthwaite's method<br>Sum Sq Mean Sq NumDF DenDF F value Pr(>F)<br>as.factor(chunk) 0.34893 0.174464 2 176 1.6346 0.1980<br>as.factor(Session) 0.02494 0.024944 1 88 0.2337 0.6300<br>as.factor(chunk):as.factor(Session) 0.28226 0.141132 2 176 1.3223 0.2692 | BH FDR Corrected p for emmeans contrasts:<br>0.5-0.1 and 1-1.5 vs. 0.0-0.5 p <.002<br>0.5-1 vs. 1-1.5 p = .11 | chunk=time bin<br>0-0.5<br>0.5-1<br>1-1.5 |
| 3J | Fisher Exact test (scipy)<br>OR = 1.84, p = .025 | BH adjusted p = .042 | |
| 3L | Mixed Effects (lmer +lmerTest)<br>Type III Analysis of Variance Table with Satterthwaite's method<br>Sum Sq Mean Sq NumDF DenDF F value Pr(>F)<br>as.factor(chunk) 0.26073 0.130367 2 134 1.2206 0.2983<br>as.factor(Session) 0.00629 0.006286 1 67 0.0589 0.8091<br>as.factor(chunk):as.factor(Session) 0.37666 0.188329 2 134 1.7633 0.1754 | | chunk=time bin<br>0-0.5<br>0.5-1<br>1-1.5 |

**Figure 4**

| Panel | Stats | Post-hoc tests/ P Adjustments | Notes |
|---|---|---|---|
| **4B** | Linear mixed model fit by REML. t-tests use Satterthwaite's method<br>Estimate Std. Error df t value Pr(>\|t\|)<br>(Intercept) 0.27682 0.03772 11.48903 7.338 1.15e-05<br>as.factor(timebin_chunk)2 -0.02697 0.02734 1247.99 -0.987 0.32403<br>as.factor(timebin_chunk)3 -0.03224 0.02734 1247.99 -1.179 0.23858<br>as.factor(timebin_chunk)4 -0.09013 0.02734 1247.99 -3.297 0.00101<br>as.factor(timebin_chunk)5 -0.12961 0.02734 1247.99 -4.740 2.38e-06<br>as.factor(Session)Psilocybin -0.08336 0.03128 1217.87 -2.665 0.00780<br>as.factor(timebin_chunk)2:as.factor(Session)Psilocybin 0.05537 0.03806 1247.99 1.455 0.14602<br>as.factor(timebin_chunk)3:as.factor(Session)Psilocybin 0.07359 0.03806 1247.99 1.933 0.05340<br>as.factor(timebin_chunk)4:as.factor(Session)Psilocybin 0.10124 0.03806 1247.99 2.660 0.00792<br>as.factor(timebin_chunk)5:as.factor(Session)Psilocybin 0.10430 0.03806 1247.99 2.740 0.00623 | BH FDR Corrected p for emmeans contrasts:<br><br>timebin_chunk contrast t.ratio p.value<br>1 Baseline - Psilocybin 2.664 0.0392<br>2 Baseline - Psilocybin 0.894 0.7097<br>3 Baseline - Psilocybin 0.312 0.7551<br>4 Baseline - Psilocybin -0.571 0.7097<br>5 Baseline - Psilocybin -0.669 0.7097 | Using lme4 and emmeans<br>Model Structure<br>value ~ as.factor(timebin_chunk) * as.factor(Session)<br>+ (1 \| id) + (1 \| id:unit)<br>id=subject, unit = unitID<br>timebin_chunk = 0.5 sec timebin average<br>e.g. -1-0.5,-0.5-0, etc |
| **4D** | Linear mixed model fit by REML. t-tests use Satterthwaite's method<br>Estimate Std. Error df t value Pr(>\|t\|)<br>(Intercept) 6.128e-02 2.531e-02 4.867e+01 2.422 0.0192<br>as.factor(timebin_chunk)2 8.553e-03 3.122e-02 1.248e+03 0.274 0.7842<br>as.factor(timebin_chunk)3 3.336e-01 3.122e-02 1.248e+03 10.685 <2e-16<br>as.factor(timebin_chunk)4 4.184e-01 3.122e-02 1.248e+03 13.403 <2e-16<br>as.factor(timebin_chunk)5 3.237e-01 3.122e-02 1.248e+03 10.369 <2e-16<br>as.factor(Session)Psilocybin 1.252e-02 3.297e-02 1.459e+03 0.380 0.7041<br>as.factor(timebin_chunk)2:as.factor(Session)Psilocybin -1.658e-02 4.346e-02 1.248e+03 -0.381 0.7030<br>as.factor(timebin_chunk)3:as.factor(Session)Psilocybin -6.195e-02 4.346e-02 1.248e+03 -1.425 0.1543<br>as.factor(timebin_chunk)4:as.factor(Session)Psilocybin -7.583e-02 4.346e-02 1.248e+03 -1.745 0.0813<br>as.factor(timebin_chunk)5:as.factor(Session)Psilocybin 2.693e-02 4.346e-02 1.248e+03 0.620 0.5356 | No interaction contrasts | Using lme4 and emmeans<br><br>Model Structure<br>value ~ as.factor(timebin_chunk) * as.factor(Session)<br>+ (1 \| id) + (1 \| id:unit)<br>id=subject, unit = unitID<br>timebin_chunk = 0.5 sec timebin average<br>e.g. -1-0.5,-0.5-0, etc |
| **4E** | Linear mixed model fit by REML. t-tests use Satterthwaite's method<br>Estimate Std. Error df t value Pr(>\|t\|)<br>(Intercept) 6.974e-02 1.414e-02 1.464e+03 4.933 9.02e-07<br>as.factor(timebin_chunk)2 1.316e-03 1.867e-02 1.248e+03 0.070 0.94382<br>as.factor(timebin_chunk)3 4.342e-02 1.867e-02 1.248e+03 2.326 0.02019<br>as.factor(timebin_chunk)4 9.276e-02 1.867e-02 1.248e+03 4.969 7.68e-07<br>as.factor(timebin_chunk)5 4.671e-02 1.867e-02 1.248e+03 2.502 0.01248<br>as.factor(Session)Psilocybin 5.572e-03 1.968e-02 1.464e+03 0.283 0.77714<br>as.factor(timebin_chunk)2:as.factor(Session)Psilocybin -2.045e-02 2.599e-02 1.248e+03 -0.787 0.43152<br>as.factor(timebin_chunk)3:as.factor(Session)Psilocybin -1.070e-02 2.599e-02 1.248e+03 -0.412 0.68051<br>as.factor(timebin_chunk)4:as.factor(Session)Psilocybin -6.931e-02 2.599e-02 1.248e+03 -2.666 0.00776<br>as.factor(timebin_chunk)5:as.factor(Session)Psilocybin -9.673e-03 2.599e-02 1.248e+03 -0.372 0.70983 | BH FDR Corrected p for emmeans contrasts:<br><br>timebin_chunk contrast t.ratio p.value<br>1 Baseline - Psilocybin -0.283 0.8349<br>2 Baseline - Psilocybin 0.756 0.8349<br>3 Baseline - Psilocybin 0.261 0.8349<br>4 Baseline - Psilocybin 3.238 0.0061<br>5 Baseline - Psilocybin 0.208 0.8349 | Using lme4 and emmeans<br><br>Model Structure<br>value ~ as.factor(timebin_chunk) * as.factor(Session) + (1\|id:unit)<br>id=subject, unit = unitID<br>Dropped random effect of 1:id because it was singular.<br>same interpretation if included anyway<br><br>timebin_chunk = 0.5 sec timebin average<br>e.g. -1-0.5,-0.5-0, etc |
| **4F** | Linear mixed model fit by REML. t-tests use Satterthwaite's method<br>Estimate Std. Error df t value Pr(>\|t\|)<br>(Intercept) 1.210e-01 2.086e-02 1.799e+01 5.800 1.71e-05<br>as.factor(timebin_chunk)2 -3.947e-03 1.877e-02 1.248e+03 -0.210 0.8335<br>as.factor(timebin_chunk)3 -3.289e-03 1.877e-02 1.248e+03 -0.175 0.8609<br>as.factor(timebin_chunk)4 -3.421e-02 1.877e-02 1.248e+03 -1.823 0.0686<br>as.factor(timebin_chunk)5 -3.553e-02 1.877e-02 1.248e+03 -1.893 0.0586<br>as.factor(Session)Psilocybin -4.913e-03 2.034e-02 1.383e+03 -0.242 0.8092<br>as.factor(timebin_chunk)2:as.factor(Session)Psilocybin 1.259e-02 2.613e-02 1.248e+03 0.482 0.6300<br>as.factor(timebin_chunk)3:as.factor(Session)Psilocybin 1.810e-02 2.613e-02 1.248e+03 0.693 0.4885<br>as.factor(timebin_chunk)4:as.factor(Session)Psilocybin 5.211e-02 2.613e-02 1.248e+03 1.994 0.0463<br>as.factor(timebin_chunk)5:as.factor(Session)Psilocybin 2.874e-02 2.613e-02 1.248e+03 1.100 0.2717 | BH FDR Corrected p for emmeans contrasts:<br><br>timebin_chunk contrast t.ratio p.value<br>1 Baseline - Psilocybin 0.241 0.8093<br>2 Baseline - Psilocybin -0.377 0.8093<br>3 Baseline - Psilocybin -0.648 0.8093<br>4 Baseline - Psilocybin -2.319 0.1026<br>5 Baseline - Psilocybin -1.171 0.6049 | Using lme4 and emmeans<br><br>Model Structure<br>value ~ as.factor(timebin_chunk) * as.factor(Session)<br>+ (1 \| id) + (1 \| id:unit)<br>id=subject, unit = unitID<br>timebin_chunk = 0.5 sec timebin average<br>e.g. -1-0.5,-0.5-0, etc |

**Figure 5**

<table>
<tr><th>Panel</th><th colspan="6">Stats</th><th>Post-hoc tests/ P Adjustments</th><th>Notes</th></tr>
<tr><td rowspan="11">5C</td><td colspan="6">Linear mixed model fit by REML. t-tests use Satterthwaite's method</td><td></td><td></td></tr>
<tr><td></td><td>Estimate</td><td>Std. Error</td><td>df</td><td>t value</td><td>Pr(>|t|)</td><td>BH FDR Corrected p for emmeans contrasts:</td><td>Using lme4 and emmeans</td></tr>
<tr><td>(Intercept)</td><td>9.228e-02</td><td>2.112e-02</td><td>1.118e+02</td><td>4.369</td><td>2.8e-05</td><td></td><td></td></tr>
<tr><td>as.factor(timebin_chunk)2</td><td>2.347e-02</td><td>2.544e-02</td><td>8.240e+02</td><td>0.923</td><td>0.35647</td><td>timebin_chunk contrast t.ratio p.value</td><td></td></tr>
<tr><td>as.factor(timebin_chunk)3</td><td>7.755e-02</td><td>2.544e-02</td><td>8.240e+02</td><td>3.049</td><td>0.00237</td><td>1 Psilocybin - Baseline 0.869 0.6576</td><td></td></tr>
<tr><td>as.factor(timebin_chunk)4</td><td>2.959e-02</td><td>2.544e-02</td><td>8.240e+02</td><td>1.163</td><td>0.24503</td><td>2 Psilocybin - Baseline 0.589 0.6948</td><td></td></tr>
<tr><td>as.factor(timebin_chunk)5</td><td>1.633e-02</td><td>2.544e-02</td><td>8.240e+02</td><td>0.642</td><td>0.52116</td><td>3 Psilocybin - Baseline -2.221 0.1331</td><td>Model Structure</td></tr>
<tr><td>as.factor(Session)Psilocy</td><td>2.488e-02</td><td>2.844e-02</td><td>8.214e+02</td><td>0.875</td><td>0.38184</td><td>4 Psilocybin - Baseline 0.852 0.6576</td><td>value ~ as.factor(timebin_chunk) * as.factor(Session)</td></tr>
<tr><td>as.factor(timebin_chunk)2:as.factor(Session)Psilocy</td><td>-8.015e-03</td><td>3.498e-02</td><td>8.240e+02</td><td>-0.229</td><td>0.81882</td><td>5 Psilocybin - Baseline 0.331 0.7409</td><td>+ (1 | id) + (1 | id:unit)</td></tr>
<tr><td>as.factor(timebin_chunk)3:as.factor(Session)Psilocy</td><td>-8.846e-02</td><td>3.498e-02</td><td>8.240e+02</td><td>-2.529</td><td>0.01163</td><td></td><td></td></tr>
<tr><td>as.factor(timebin_chunk)4:as.factor(Session)Psilocy</td><td>-5.009e-04</td><td>3.498e-02</td><td>8.240e+02</td><td>-0.014</td><td>0.98858</td><td></td><td>id=subject, unit = unitID</td></tr>
<tr><td></td><td>as.factor(timebin_chunk)5:as.factor(Session)Psilocy</td><td>-1.542e-02</td><td>3.498e-02</td><td>8.240e+02</td><td>-0.441</td><td>0.65950</td><td></td><td></td></tr>
<tr><td rowspan="12">5F</td><td colspan="6">Linear mixed model fit by REML. t-tests use Satterthwaite's method</td><td></td><td></td></tr>
<tr><td></td><td>Estimate</td><td>Std. Error</td><td>df</td><td>t value</td><td>Pr(>|t|)</td><td></td><td>Using lme4 and emmeans</td></tr>
<tr><td>(Intercept)</td><td>0.088277</td><td>0.025214</td><td>18.184632</td><td>3.501</td><td>0.00252</td><td>No interaction contrasts</td><td></td></tr>
<tr><td>as.factor(timebin_chunk)2</td><td>-0.016327</td><td>0.021552</td><td>823.998647</td><td>-0.758</td><td>0.44894</td><td></td><td>Model Structure</td></tr>
<tr><td>as.factor(timebin_chunk)3</td><td>0.030612</td><td>0.021552</td><td>823.998647</td><td>1.420</td><td>0.15587</td><td></td><td>value ~ as.factor(timebin_chunk) * as.factor(Session)</td></tr>
<tr><td>as.factor(timebin_chunk)4</td><td>0.007143</td><td>0.021552</td><td>823.998647</td><td>0.331</td><td>0.74041</td><td></td><td>+ (1 | id) + (1 | id:unit)</td></tr>
<tr><td>as.factor(timebin_chunk)5</td><td>-0.009184</td><td>0.021552</td><td>823.998647</td><td>-0.426</td><td>0.67014</td><td></td><td></td></tr>
<tr><td>as.factor(Session)Psilocy</td><td>0.006549</td><td>0.025158</td><td>747.398452</td><td>0.260</td><td>0.79469</td><td></td><td>id=subject, unit = unitID</td></tr>
<tr><td>as.factor(timebin_chunk)2:as.factor(Session)Psilocy</td><td>0.019054</td><td>0.029636</td><td>823.998647</td><td>0.643</td><td>0.52046</td><td></td><td></td></tr>
<tr><td>as.factor(timebin_chunk)3:as.factor(Session)Psilocy</td><td>-0.016067</td><td>0.029636</td><td>823.998647</td><td>-0.542</td><td>0.58788</td><td></td><td></td></tr>
<tr><td>as.factor(timebin_chunk)4:as.factor(Session)Psilocy</td><td>0.006494</td><td>0.029636</td><td>823.998646</td><td>0.219</td><td>0.82662</td><td></td><td></td></tr>
<tr><td>as.factor(timebin_chunk)5:as.factor(Session)Psilocy</td><td>-0.008998</td><td>0.029636</td><td>823.998646</td><td>-0.304</td><td>0.76150</td><td></td><td></td></tr>
</table>

**Figure 6**

<table>
<tr><th>Panel</th><th>Statistics</th><th>Post-hoc tests/ P Adjustments</th><th>Notes</th></tr>
<tr><td>6A (Baseline)</td><td>Sum Sq Df F value Pr(>F)<br>(Intercept) 8.1409 1 6373.282 < 2.2e-16 ***<br>size 1.3283 5 207.972 < 2.2e-16 ***<br>Session 0.2920 1 228.586 < 2.2e-16 ***<br>cluster:Session 0.5425 5 84.946 < 2.2e-16 ***<br>Residuals 0.2912 228</td><td>Observed vs shuffle t ratio p value<br>10 15.1 <.001<br>50 27.4 <.001<br>100 33.1 <.001<br>150 36.4 <.001<br>200 38.6 <.001<br>Full 39.4 <.001</td><td>Used car and emmeans<br>Session is shuffle vs observed here<br>size is pseudopopulation size</td></tr>
<tr><td>6A (Psilocybin)</td><td>Sum Sq Df F value Pr(>F)<br>(Intercept) 5.8699 1 3979.545 < 2.2e-16 ***<br>size 1.1531 5 156.353 < 2.2e-16 ***<br>Session 0.0595 1 40.327 1.15e-09 ***<br>cluster:Session 0.4074 5 55.236 < 2.2e-16 ***<br>Residuals 0.3363 228</td><td>Observed vs shuffle t ratio p value<br>10 6.4 <.001<br>50 12.3 <.001<br>100 18 <.001<br>150 21.8 <.001<br>200 23.6 <.001<br>Full 25.9 <.001</td><td>Used car and emmeans<br>Session is shuffle vs observed here<br>size is pseudopopulation size</td></tr>
<tr><td>6A</td><td>Sum Sq Df F value Pr(>F)<br>(Intercept) 8.1409 1 5169.5145 < 2.2e-16 ***<br>size 1.3283 5 168.6909 < 2.2e-16 ***<br>Session 0.0926 1 58.8274 4.958e-13 ***<br>cluster:Session 0.0118 5 1.4994 0.1909<br>Residuals 0.3591 228</td><td>contrast t.ratio p.value<br>cluster50 - cluster10 14.404 <.0001<br>cluster100 - cluster10 22.518 <.0001<br>cluster150 - cluster10 28.561 <.0001<br>cluster200 - cluster10 31.576 <.0001<br>cluster250 - cluster10 32.414 <.0001</td><td>Used car and emmeans<br><br>Session is psilocybin vs baseline here<br>size is pseudopopulation size</td></tr>
<tr><td>6C</td><td>Sum Sq Df F value Pr(>F)<br>(Intercept) 0.07175 1 24.6512 1.524e-06 ***<br>cluster 0.09172 4 7.8783 6.767e-06 ***<br>Session 0.05447 1 18.7142 2.451e-05 ***<br>cluster:Session 0.01883 4 1.6174 0.1715<br>Residuals 0.55302 190</td><td>contrast t.ratio p.value<br>R_T2 - ActionDecay 2.893 0.0043<br>R_TI - ActionDecay 7.616 <.0001<br>NR_T1 - ActionDecay 6.116 <.0001<br>NR_T2 - ActionDecay 4.246 <.0001</td><td>Used car and emmeans<br><br>Session is psilocybin vs baseline here<br>Cluster is subcluster type from Fig 3.<br>e.g. R_T1= Reward Type I</td></tr>
</table>

**Supplemental Data 1**

<table>
<tr><th>Panel</th><th>Stats</th><th>Post-hoc tests/ P Adjustments</th><th>Notes</th></tr>
<tr><td>1B</td><td>statsmodels RM anova<br><br>F Value Num DF Den DF Pr > F<br>Session 6.6709 1.0000 7.0000 0.0363<br>priorR 1.1850 1.0000 7.0000 0.3124<br>Session:priorR 0.0020 1.0000 7.0000 0.9657</td><td></td><td></td></tr>
<tr><td>1C</td><td>statsmodels RM anova<br><br>F Value Num DF Den DF Pr > F<br>Session 15.4750 1.0000 7.0000 0.0056<br>outcome 34.0238 1.0000 7.0000 0.0006<br>Session:outcome 10.4699 1.0000 7.0000 0.0143</td><td>post hoc t tests (baseline vs psilocybin)<br><br>Rewarded trials : t(7)=-1.4, p=.20<br>No reward trials: t(7)=-4.05, p= .0049</td><td></td></tr>
<tr><td>1D</td><td>paired t test<br>T(7)=0.63 P = 0.54</td><td></td><td></td></tr>
<tr><td>1E</td><td>Generalized linear mixed model fit by maximum likelihood<br>Estimate Std. Error z value Pr(>|z|)<br>(Intercept) 0.50410 0.24929 2.022 0.0432<br>SessionPsilocybin 0.77036 0.36374 2.118 0.0342<br>phase -0.62861 0.28627 -2.196 0.0281<br>post0 0.04608 0.04924 0.936 0.3493<br>switchtype 0.29561 0.28354 1.043 0.2972<br>SessionPsilocybin:phase -0.21933 0.48735 -0.450 0.6527<br>SessionPsilocybin:post0 -0.09251 0.06904 -1.340 0.1803<br>post0:switchtype -0.09146 0.05098 -1.794 0.0728</td><td></td><td>Models fit on raw choice data<br>so binomial (logit) family was used<br><br>Switch type captures low-high vs low-low effect</td></tr>
<tr><td>1F</td><td>Generalized linear mixed model fit by maximum likelihood<br>Estimate Std. Error z value Pr(>|z|)<br>(Intercept) 0.22812 0.34089 0.669 0.503<br>SessionPsilocybin 0.04749 0.55606 0.085 0.932<br>phase -0.13910 0.41416 -0.336 0.737<br>post0 0.05235 0.06221 0.842 0.400<br>SessionPsilocybin:phase -0.38710 0.67673 -0.572 0.567<br>SessionPsilocybin:post0 0.05346 0.10185 0.525 0.600</td><td></td><td>Models fit on raw choice data<br>so binomial (logit) family was used<br><br>1 rat excluded (only 2 high uncertainty trials)</td></tr>
<tr><td>1G</td><td>Generalized linear mixed model fit by maximum likelihood<br><br>Estimate Std. Error z value Pr(>|z|)<br>(Intercept) 0.300323 0.222694 1.349 0.177<br>SessionSaline 0.293920 0.318898 0.922 0.357<br>phase -0.266294 0.330879 -0.805 0.421<br>post0 0.027893 0.049299 0.566 0.572<br>SessionSaline:phase -0.304328 0.470898 -0.646 0.518<br>SessionSaline:post0 0.002573 0.069790 0.037 0.971</td><td></td><td></td></tr>
<tr><td>1G (right)</td><td>paired t-test<br>t(3)=-1.21, P=0.31</td><td></td><td>scipy ttest_rel<br>1 rat excluded<br>Saline: only 2 post transtions trials/only 20 trials total</td></tr>
</table>

**Supplemental Data 3**

| Panel | Stats | | | Notes |
|---|---|---|---|---|
| **1C** | unpaired t-test | t(427)=0.20 | p=.83 | scipy ttest_ind |

**Supplemental Data 5**

| Panel | Stats | Post-hoc tests/ P Adjustments | Notes |
|---|---|---|---|
| 1C | Linear mixed model fit by REML. t-tests use Satterthwaite's method<br>Estimate Std. Error df t value Pr(>\|t\|)<br>(Intercept) 3.580e-02 1.234e-02 1.108e+02 2.902 0.00448<br>as.factor(timebin_chunk)2 4.539e-02 1.517e-02 1.248e+03 2.993 0.00282<br>as.factor(timebin_chunk)3 3.684e-02 1.517e-02 1.248e+03 2.429 0.01527<br>as.factor(timebin_chunk)4 1.382e-02 1.517e-02 1.248e+03 0.911 0.36249<br>as.factor(timebin_chunk)5 3.290e-02 1.517e-02 1.248e+03 2.169 0.03027<br>as.factor(Session)Psilocybin 3.740e-02 1.614e-02 1.441e+03 2.318 0.02061<br>as.factor(timebin_chunk)2:as.factor(Session)Psilocybin -3.675e-02 2.111e-02 1.248e+03 -1.741 0.08199<br>as.factor(timebin_chunk)3:as.factor(Session)Psilocybin -4.301e-02 2.111e-02 1.248e+03 -2.037 0.04184<br>as.factor(timebin_chunk)4:as.factor(Session)Psilocybin -7.026e-03 2.111e-02 1.248e+03 -0.333 0.73939<br>as.factor(timebin_chunk)5:as.factor(Session)Psilocybin -1.252e-02 2.111e-02 1.248e+03 -0.593 0.55318 | timebin_chunk contrast t.ratio p.value<br>1 Baseline - Psilocybin -2.314 0.1041<br>2 Baseline - Psilocybin -0.040 0.9682<br>3 Baseline - Psilocybin 0.347 0.9103<br>4 Baseline - Psilocybin -1.879 0.1511<br>5 Baseline - Psilocybin -1.539 0.2068 | |
| 1D | Linear mixed model fit by REML. t-tests use Satterthwaite's method<br>Estimate Std. Error df t value Pr(>\|t\|)<br>(Intercept) 6.699e-02 9.996e-03 1.839e+02 6.701 2.44e-10<br>as.factor(timebin_chunk)2 9.868e-03 1.311e-02 1.248e+03 0.753 0.452<br>as.factor(timebin_chunk)3 -6.579e-03 1.311e-02 1.248e+03 -0.502 0.616<br>as.factor(timebin_chunk)4 1.908e-02 1.311e-02 1.248e+03 1.455 0.146<br>as.factor(timebin_chunk)5 5.921e-03 1.311e-02 1.248e+03 0.452 0.652<br>as.factor(Session)Psilocybin -1.363e-02 1.369e-02 1.487e+03 -0.996 0.319<br>as.factor(timebin_chunk)2:as.factor(Session)Psilocybin -2.283e-02 1.826e-02 1.248e+03 -1.251 0.211<br>as.factor(timebin_chunk)3:as.factor(Session)Psilocybin 1.646e-02 1.826e-02 1.248e+03 0.901 0.368<br>as.factor(timebin_chunk)4:as.factor(Session)Psilocybin -2.649e-02 1.826e-02 1.248e+03 -1.451 0.147<br>as.factor(timebin_chunk)5:as.factor(Session)Psilocybin -2.320e-02 1.826e-02 1.248e+03 -1.271 0.204 | | |
| 1E | Linear mixed model fit by REML. t-tests use Satterthwaite's method<br>Estimate Std. Error df t value Pr(>\|t\|)<br>(Intercept) 1.210e-01 2.086e-02 1.799e+01 5.800 1.71e-05<br>as.factor(timebin_chunk)2 -3.947e-03 1.877e-02 1.248e+03 -0.210 0.8335<br>as.factor(timebin_chunk)3 -3.289e-03 1.877e-02 1.248e+03 -0.175 0.8609<br>as.factor(timebin_chunk)4 -3.421e-02 1.877e-02 1.248e+03 -1.823 0.0686<br>as.factor(timebin_chunk)5 -3.553e-02 1.877e-02 1.248e+03 -1.893 0.0586<br>as.factor(Session)Psilocybin -4.913e-03 2.034e-02 1.383e+03 -0.242 0.8092<br>as.factor(timebin_chunk)2:as.factor(Session)Psilocybin 1.259e-02 2.613e-02 1.248e+03 0.482 0.6300<br>as.factor(timebin_chunk)3:as.factor(Session)Psilocybin 1.810e-02 2.613e-02 1.248e+03 0.693 0.4885<br>as.factor(timebin_chunk)4:as.factor(Session)Psilocybin 5.211e-02 2.613e-02 1.248e+03 1.994 0.0463<br>as.factor(timebin_chunk)5:as.factor(Session)Psilocybin 2.874e-02 2.613e-02 1.248e+03 1.100 0.2717 | timebin_chunk contrast t.ratio p.value<br>1 Baseline - Psilocybin 0.241 0.8093<br>2 Baseline - Psilocybin -0.377 0.8093<br>3 Baseline - Psilocybin -0.648 0.8093<br>4 Baseline - Psilocybin -2.319 0.1026<br>5 Baseline - Psilocybin -1.171 0.6049 | |

Supplemental Data 7

| Panel | Stats | Post-hoc tests/ P Adjustments | Notes |
|---|---|---|---|
| 1A | Sum Sq Df F value Pr(>F)<br>(Intercept) 5.7800 1 6737.0697 <2e-16<br>size 0.8210 5 191.3828 <2e-16<br>Session 0.0013 1 1.5093 0.2205<br>cluster:Session 0.0035 5 0.8131 0.5414<br>Residuals 0.1956 228 | | Used car<br>Session is psilocybin vs baseline here<br>size is pseudopopulation size |
| 1B (Baseline) | Sum Sq Df F value Pr(>F)<br>(Intercept) 9.7580 1 4979.653 < 2.2e-16<br>size 1.3368 5 136.441 < 2.2e-16<br>Session 0.4951 1 252.637 < 2.2e-16<br>cluster:Session 0.5408 5 55.197 < 2.2e-16<br>Residuals 0.4468 228 | | Used car and<br>Session is shuffle vs observed here<br>size is pseudopopulation size |
| 1B (Psilocybin) | Sum Sq Df F value Pr(>F)<br>(Intercept) 8.5543 1 3977.841 < 2.2e-16<br>size 1.6988 5 157.988 < 2.2e-16<br>Session 0.4020 1 186.935 < 2.2e-16<br>cluster:Session 0.6501 5 60.462 < 2.2e-16<br>Residuals 0.4903 228 | | Used car<br><br>Session is psilocybin vs baseline here<br>size is pseudopopulation size |
| 1B | Sum Sq Df F value Pr(>F)<br>(Intercept) 9.7580 1 6046.7928 < 2.2e-16<br>size 1.3368 5 165.6807 < 2.2e-16<br>Session 0.0198 1 12.2711 0.0005535<br>cluster:Session 0.0261 5 3.2374 0.0076263<br>Residuals 0.3679 228 | FDR-BH corrected<br>Baseline vs Psilocybin<br>t p value<br>10 3.5 0.002<br>50 5.49 <.0001<br>100 2.49 0.026<br>150 1.33 0.27<br>200 1.2 0.28<br>250 0.71 0.48 | Used car and emmeans for post hoc<br><br>Session is psilocybin vs baseline here<br>size is pseudopopulation size |
| 1D | Sum Sq Df F value Pr(>F)<br>(Intercept) 0.0022077 1 16.4304 7.355e-05<br>cluster 0.0041427 4 7.7078 8.914e-06<br>Session 0.0072091 1 53.6525 6.626e-12<br>cluster:Session 0.0045054 4 8.3827 3.005e-06<br>Residuals 0.0255298 190 | FDR-BH corrected<br>Comparing action decay to other clusters<br>(either ns or action decay had less effect)<br>Session contrast t.ratio p.value<br>BL R_T2 - ActionDecay -1.802 0.0974<br>BL R_TI - ActionDecay -3.513 0.0009<br>BL NR_T1 - ActionDecay -0.028 0.9777<br>BL NR_T2 - ActionDecay -4.286 0.0001<br>PS R_T2 - ActionDecay -7.926 <.0001<br>PS R_TI - ActionDecay -6.831 <.0001<br>PS NR_T1 - ActionDecay 1.213 0.2592<br>PS NR_T2 - ActionDecay -5.731 <.0001 | Used car and emmeans for posthoc<br><br>Session is psilocybin vs baseline here<br>Cluster is subcluster type from Fig 3.<br>e.g. R_T1= Reward Type I |
